# Spatial joint profiling of DNA methylome and transcriptome in mammalian tissues

**DOI:** 10.1101/2025.07.01.662607

**Authors:** Chin Nien Lee, Hongxiang Fu, Angelysia Cardilla, Wanding Zhou, Yanxiang Deng

## Abstract

The spatial resolution of omics dynamics is fundamental to understanding tissue biology. Spatial profiling of DNA methylation, which is a canonical epigenetic mark extensively implicated in transcriptional regulation, remains an unmet demand. Here, we introduce a method for whole genome spatial co-profiling of DNA methylation and transcriptome of the same tissue section at near single-cell resolution. Applying this technology to mouse embryogenesis and postnatal brain resulted in rich DNA-RNA bimodal tissue maps. These maps revealed the spatial context of known methylation biology and its interplay with gene expression. The two modalities’ concordance and distinction in spatial patterns highlighted a synergistic molecular definition of cell identity in spatial programming of mammalian development and brain function. By integrating spatial maps of mouse embryos at two different developmental stages, we reconstructed the dynamics of both epigenome and transcriptome underlying mammalian embryogenesis, revealing details in sequence, cell type, and region-specific methylation-mediated transcriptional regulation. This method extends the scope of spatial omics to DNA cytosine methylation for a more comprehensive understanding of tissue biology over development and disease.

## Main

DNA methylation, which involves cytosine modifications such as the 5’-methylation (5mC) and 5’-hydroxymethylation (5hmC), is a key epigenetic mechanism that modulates gene expression and lineage specification by regulating chromatin structure and accessibility to transcriptional machinery^1,2^. This epigenetic modification varies dynamically across different cell types, developmental stages, and environmental conditions^3^. Abnormal cytosine methylation patterns are associated with various diseases, including cancer ^4^, autoimmune diseases^5^, and inflammation^6^. In addition, altered DNA methylation levels have been observed in aging tissues^7^, contributing to biomarkers and predictive models (DNA methylation clock) for chronological and biological age^8^.

Despite advancements in single-cell methylome analysis, which provided valuable insights into epigenetic mechanisms orchestrating biological diversity^9-16^, the lack of spatial information within intact tissues limited our understanding of gene regulation during tissue development and disease progression. Emerging spatial multi-omics technologies have begun to address this gap by enabling the detection of various omics profiles within tissue microenvironments. These technologies have significantly advanced our understanding of biological complexity by allowing the study of molecular analytes in their native tissue contexts. However, current spatial methodologies are limited to mapping histone modifications and chromatin accessibility, the transcriptome, and a select panel of proteins^17-20^. Directly measuring spatial DNA methylation has not been possible, leaving this crucial epigenetic layer largely unexplored.

Here, we introduce a technology for spatial joint profiling of DNA Methylome and Transcriptome (Spatial-DMT) on the same tissue section at near single-cell resolution. We applied Spatial-DMT to profile mouse embryos and postnatal mouse brains. The spatial maps uncovered intricate spatiotemporal regulatory mechanisms of gene expression within the tissue context. Spatial-DMT investigates interactive molecular hierarchies in development, physiology, and pathogenesis in a spatially resolved manner.

### Spatial-DMT design and workflow

Spatial-DMT combines microfluidic in situ barcoding^17^, cytosine deamination conversion^16^, and high-throughput Next-Generation Sequencing to achieve spatial methylome profiling directly in tissue. The experimental workflow of Spatial-DMT was illustrated in Figures 1a and b, Extended Data Fig. 1, and a step-by-step protocol and detailed information for each reagent used in this protocol were provided in Supplementary Tables 1 to 4.

**Fig. 1:**
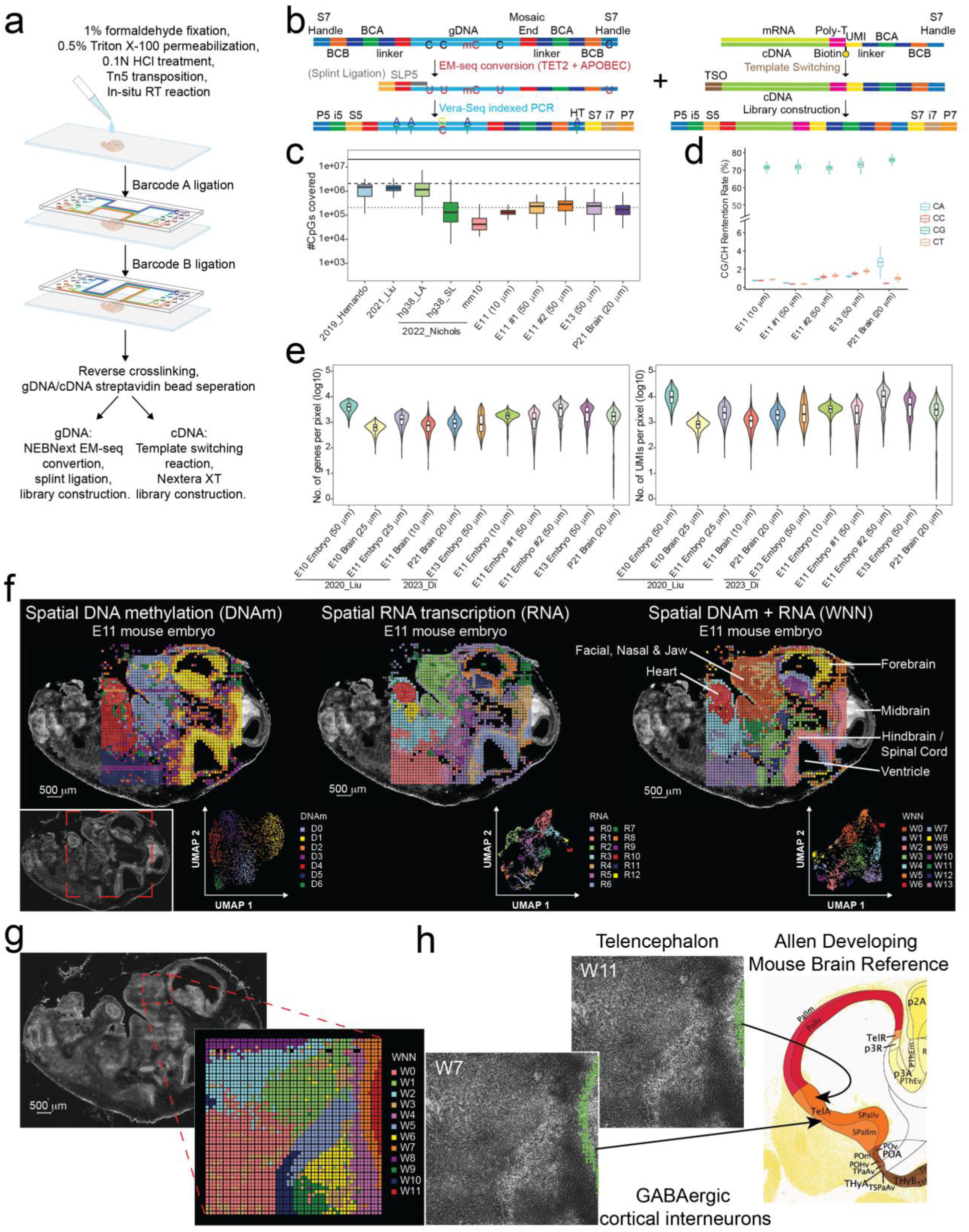
Overview of Spatial-DMT for simultaneous profiling of DNA methylation and transcriptome in tissues. **a,** Schematic diagram of the Spatial-DMT workflow. **b,** Chemistry workflow of DNA methylation and RNA library preparation. The tissue section is first fixed and permeabilized, followed by HCl treatment, Tn5 transposition, and in-situ reverse transcription (RT). Barcodes A and B are ligated sequentially. Streptavidin bead separation allows for differential processing: cDNA is subjected to template switching reaction and Nextera XT library preparation, while gDNA undergoes NEBNext EM-seq conversion, splint ligation, and library construction. **c,** Comparison of the number of CpGs covered per tissue pixel/cell between Spatial-DMT and other single-cell DNA methylation datasets^7,16,22,23^. **d,** Box plots showing CG/CH retention rates in E11 embryo, E13 embryo, and P21 brain tissues. **e,** The number of genes (left) and the number of UMIs (right) per tissue pixel for Spatial-DMT. The box plots show the median (center line), the first and third quartiles (box limits), and 1.5x the interquartile range (whiskers). **f,** Spatial distribution and UMAP visualization of DNA methylation (left), RNA transcription (middle), and integrated DNA methylation and RNA (WNN) (right) in E11 embryo (pixel size, 50 μm, ROI area 5x5 mm^2^). The red dashed box in the bright-field image indicates the region of interest for mapping. Spatial maps show distinct clustering patterns, with each color representing a different cluster. Integration of both DNA methylation and RNA transcription data reveals more refined spatial clusters and highlights specific anatomical regions such as the forebrain, midbrain, hindbrain/spinal cord, ventricle, heart, and facial, nasal & jaw regions. **g,** Spatial mapping and joint clustering of DNA methylation and RNA data in E11 facial and forebrain regions (pixel size, 10 μm, ROI area 1x1 mm^2^). The red dashed box in the bright-field image indicates the region of interest. **h,** Spatial mapping of GABAergic cortical interneurons and telencephalon cells identified by cell type decomposition of spatial transcriptomic pixels using a mouse embryo scRNA-seq reference^36^. Overlay of cell annotations with the tissue image shows that spatial clusters align with the anatomic references from the Allen Developing Mouse Atlases^84^.

Specifically, hydrochloric acid (HCl) was applied to a fixed frozen tissue section to disrupt the protein structure and remove nucleosome histones to improve Tn5 transposome accessibility for whole genome methylation profiling^21^. Next, Tn5 transposition was performed, and adapters containing a universal ligation linker were inserted into genomic DNA (gDNA). To further reduce the size of the large gDNA fragment and improve yield, we adopted a multi-tagmentation strategy as previously demonstrated^21^. Specifically, we implemented two rounds of tagmentation to balance DNA yield with experimental time and minimize the risk of RNA degradation. mRNAs were then captured by the biotinylated RT-primer (poly-dT primer with UMIs and a universal linker sequence; Supplementary Table 1), followed by reverse transcription to synthesize the complementary DNA (cDNA). Both genomic fragments and cDNA in the tissue were then ligated to spatial barcodes A and B sequentially. In brief, two sets of spatial barcodes (barcodes A1-A50 and B1-B50, Supplementary Tables 2 and 3) flowed perpendicularly to each other in the microfluidic channels. They were covalently conjugated to the universal linker via the templated ligation. This results in a two-dimensional grid of spatially barcoded tissue pixels within the region of interest (ROI), each defined by the unique combination of barcodes Ai and Bj (i=1-50, j=1-50; barcoded pixels, n=2,500). Barcoded gDNA fragments and cDNA were then released after reverse crosslinking. Afterward, the biotin-labeled cDNA was enriched by the streptavidin beads and separated from the gDNA-contained supernatant. cDNA was followed by template switching reaction and the cDNA library construction (Fig. 1b (right) and Extended Data Fig. 1). gDNA was treated with Enzymatic Methyl-seq conversion, splint ligation, and DNA library construction (Fig. 1b (left), Extended Data Fig. 1 and see Methods).

To minimize DNA damage during methylome profiling, we employed Enzymatic Methyl-seq, an enzyme-based alternative to bisulfite conversion (Fig. 1b and Extended Data Fig. 1). In this process, modified cytosines (the sum of 5mC and 5hmC) were both oxidized by the Ten-Eleven Translocation Methycytosine Dioxygenase 2 (TET2) protein and protected from deamination by the APOBEC protein, while unmodified cytosines were deaminated to uracil (U). Following the conversion, the SLP5 adapter with 10 random H (A, C, or T) nucleotides was ligated to the 3’ end of gDNA, adding the other PCR handle^16^. The gDNA fragments were then amplified using uracil-literate VeraSeq Ultra DNA polymerase. Since residue Cs on the PCR handle of barcode B was changed to residue Ts after deamination and PCR, we modified the P7 primer (N70X-HT) by replacing residue C with T to complement residue A on the PCR handle and amplify the fragments. Barcodes have been designed such that no cross-talk under C-to-T deamination is possible.

### Spatial-DMT yields high-quality maps of DNA methylomes and transcriptomes

To demonstrate the capability of co-profiling DNA methylation and transcription in complex tissues, we performed Spatial-DMT on embryonic day 11 (E11) and day 13 (E13) mouse embryos, and the postnatal day 21 (P21) mouse brain. Mouse embryos were profiled both at 10 μm and 50 μm pixel resolutions, while the mouse brains were profiled at a resolution of 20 μm. We developed a computational pipeline to facilitate data processing and downstream analyses (Extended Data Fig. 2). Averaging across pixels, the replicate maps from two E11 embryos with similar body positions exhibited high correlations for DNA methylation across Variably Methylated Regions (VMRs, Methods) (Pearson’s r = 0.9836) and RNA expression across genes (Pearson’s r = 0.9752), suggesting high reproducibility of our method (Extended Data Fig. 3a). Integration of the two replicate spatial maps revealed co-localization of pixels from both datasets in the UMAP co-embedding (Extended Data Fig 3b). Pixels grouped within the same UMAP clusters are consistently mapped to similar anatomical and spatial locations (Extended Data Fig. 3c). Notably, genes associated with distinct anatomical structures, including *Frem1* (facial), *Ank3* (brain), and *Trim55* (heart), exhibited strong expression and colocalization within their respective tissue contexts (Extended Data Fig. 3d-f).

For DNA methylation analysis, a total of 2,756,840,516 (E11 embryo, 10 μm), 2,864,519,160 (E11#1 embryo, 50 μm), 2,863,772,201 (E11#2 embryo, 50 μm), 3,018,600,847 (E13 embryo, 50 μm), and 3,914,623,812 (P21 brain, 20 μm) raw sequencing reads were generated (Supplementary Table 5). Low-quality pixels were filtered based on the knee-plot cutoff threshold (Extended Data Fig. 4a). After stringent quality control, 32.2%, 65.72%, 61.89%, 55.69%, and 42.48% of reads (887,671,712; 1,882,630,968; 1,772,395,274; 1,680,936,562; 1,662,805,699) were retained, yielding 355,069, 753,052, 708,958, 672,375, and 665,122 reads per pixel across 2,493, 1,954, 1,947, 1,699, and 2,235 pixels in the E11, E13, and P21 samples, respectively (Supplementary Table 5). The average number of CpG covered per pixel was approximately 136,639 (E11 embryo, 10 μm), 238,040 (E11#1 embryo, 50 μm), 281,447 (E11#2 embryo, 50 μm), 255,402 (E13 embryo, 50 μm), and 194,224 (P21 brain, 20 μm) (Supplementary Table 5), comparable to previous single-cell DNA methylation studies of mouse embryos and brain samples^7,16,22,23^ (Fig. 1c). Duplication rates ranged from approximately 20% to 53% across samples (Extended Data Fig. 4b). The retention rate, defined as the percentage of unconverted cytosines due to methylation or incomplete conversion^16,24^, of mitochondrial DNA was minimal (below 1%) (Extended Data Fig. 4c). The CpG retention rates were consistently between 70% to 80% across all samples, while the methylation level of non-CpG sites (mCH; H denotes A, C, or T) remained low (mCA <1% in embryos and about 3% to 4% in the postnatal brain) (Fig. 1d). Notably, a higher level of mCA was observed in the mouse brain, consistent with the known enrichment of non-CpG methylation in neuronal tissues^22,25^. Spatial mapping of global methylation levels revealed higher mCG levels in the spinal cord of embryos, as well as higher mCG and mCA levels in the cortex of the P21 brain (Extended Data Fig. 4d). Analysis of the methylation-free linker sequences showed that over 99% of cytosines were successfully converted to uracil, further confirming efficient enzymatic conversion (Extended Data Fig. 4e). Additionally, no evidence of RNA contamination, such as Poly A, Poly T, or TSO sequences, was detected in the DNA methylation libraries (Extended Data Fig. 4f).

We further assessed the genomic distribution of CpG coverage and found it to be uniformly distributed across genomic regions (Extended Data Fig. 5a), and methylation levels across various chromatin states were consistent with known biology and were comparable with those published databases^7,16,22,23,26^ (Extended Data Fig. 5b). For instance, DNA methylation levels were low at transcription start sites (TSS) but increased upstream and downstream of these regions (Extended Data Fig. 5c). Collectively, our approach yielded accurate and unbiased profiling of DNA methylation across the genome.

Simultaneously, we generated high-quality RNA data from the same tissue sections, enabling direct comparisons between transcriptional activity and epigenetic states. Specifically, we identified expression from 23,822 to 28,695 genes in our spatial maps (Supplementary Table 5). At the pixel level, the average number of detected genes per pixel ranges from 1,890 (E11 embryo, 10 μm; 3,596 UMIs) to 4,626 (E11 embryo, 50 μm; 16,709 UMIs), comparable to previous spatial transcriptomic studies of the mouse embryo and brain samples^17,18^ (Fig. 1e and Supplementary Table 5). Lower-resolution pixels (e.g., 50 μm) captured more UMIs and expressed genes, likely due to the inclusion of more cells within each pixel.

### Spatial co-profiling of DNA methylome and transcriptome in mouse embryo

Mammalian embryogenesis is an intricately programmed process with complex gene expression dynamics regulated by epigenetic mechanisms, including DNA methylation, at spatiotemporal scales^27^. We applied Spatial-DMT to produce spatial tissue maps of the DNA methylation and gene expression of the E11 mouse embryo. Unsupervised clustering was performed on DNA methylation and RNA transcription data (Fig. 1f). The DNA methylome and transcriptome define cell identity independently, with pixels clustered by methylation readouts sampled from variably methylated regions (VMRs)^28^ and expression levels from variably transcribed genes (Fig. 1f (left, middle) and Extended Data Fig. 5d). The two modalities can be integrated to achieve improved discrimination of inter-cellular and spatial diversity using a weighted nearest neighbor (WNN) method^29^, leading to definition of 14 clusters (Fig. 1f (right)). Superimposing the WNN clusters with histological images suggests clear correspondence to anatomical structures, e.g. craniofacial region (W0), hindbrain/spinal cord (W2), and embryonic heart (W6), consistent with the tissue histology (Fig. 1f (lower left)).

The correlations between single-modality (RNA and DNA methylation) clusters and their integrated WNN clustering results were evaluated (Fig. 1f; Extended Data Fig. 5e, f). Each modality captured distinct yet complementary aspects of cellular identity, and their integration via WNN analysis yielded clusters with enhanced resolution. RNA expression profiles exhibited a broader dynamic range, which may result in distinct clustering granularity across modalities. To quantify the relative contributions of each modality to the integrated clusters, we computed modality weights for individual spatial pixels (Extended Data Fig. 5g). This analysis revealed that some clusters were predominantly driven by gene expression (e.g., W6, cardiac tissue), while others were strongly influenced by DNA methylation (e.g., W11, craniofacial region). Collectively, these results highlight the advantage of spatial multi-omics integration in resolving cellular heterogeneity not captured by single modalities.

We examined VMRs methylations and the expression levels of neighboring genes, identified as differentially specific to the brain (W2), craniofacial (W0), and the heart (W6) regions of Figure 1f (Fig. 2a, b -cluster W2; Extended Data Fig. 7a, b -cluster W0; Extended Data Fig. 8a, b -cluster W6). Spatial cluster-specific gene expressions are frequently associated with low DNA methylation at the VMRs neighboring these genes, as exemplified by signature gene *Runx2*, *Mapt*, *Trim55* marking the craniofacial parts (jaw/upper nasal), the brain/spinal cord, and the heart, respectively (Fig. 2c; see Extended Data Figs. 6, 7-8c, for additional signature gene examples). Testing each cluster’s epigenetically regulated genes for Gene Ontology (GO) enrichment identified the developmental process specific to the anatomical structures (Extended Data Fig. 9a). While the canonical negative correlations between DNA methylation and RNA expression have been observed in many genes, we also identified genes, e.g., *Ank3*, *Atp11c*, *Cyfip2*, *Lmin*, and *Khdrbs2*, whose expression is positively correlated with the methylation levels of the associated VMRs (Fig. 2d). For instance, *Ank3*, primarily located at the axonal initial segment and nodes of Ranvier within neurons of the central and peripheral nervous systems^30^, exhibited high expression levels and DNA methylation in the brain region (Fig. 2c). The positive correlations of DNA methylation to RNA expression have been documented at enhancers^31^, gene bodies^32^, and Polycomb targets^33,34^, suggesting a complex mechanism of DNA methylation in transcriptional regulation, contingent on its interaction with transcriptional factor (TF) binding and histone modifications.

**Fig. 2:**
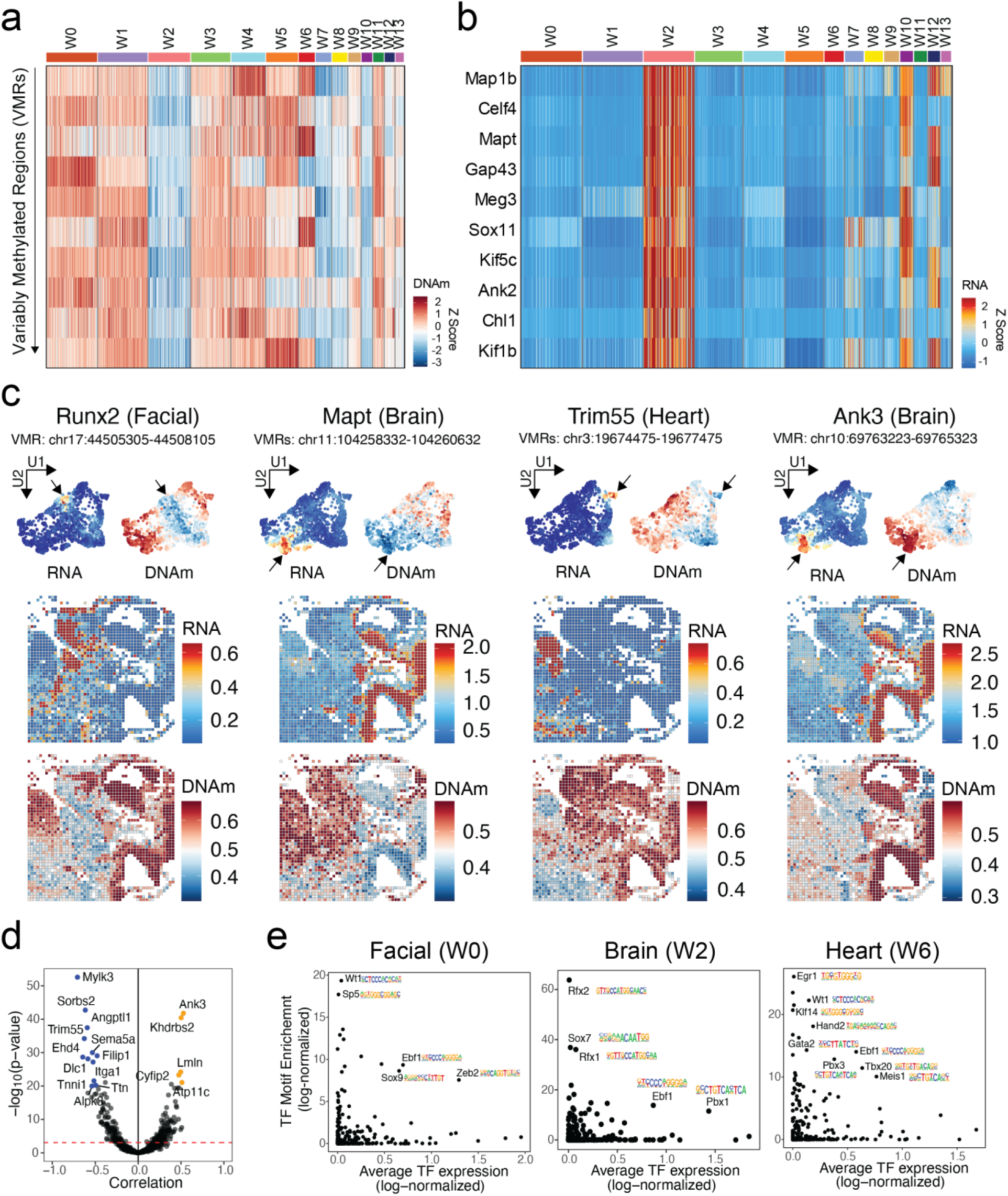
Analysis of DNA methylation and gene expression in E11 embryo. **a,** Heatmap of DNA methylation levels for the top 10 differentially methylated genomic loci in brain and spinal cord (cluster W2) of E11 embryo. Each row represents a specific genomic locus, and each column represents a different cluster. The color scale indicates the Z-scores of DNA methylation levels. **b,** Heatmap of expression levels for nearby genes corresponding to the genomic loci in **a**. Each row represents a specific gene, and each column represents a different cluster. The color scale indicates the Z-scores of gene expression levels. **c,** UMAP visualization and spatial mapping of DNA methylation (methylation percentage) and RNA expression levels (log normalized expression) for selected marker genes across different clusters. **d,** Correlation analysis between DNA methylation and RNA expression of nearby genes. Negative correlations (blue) indicate a repressive effect of methylation, while positive correlations (orange) indicate activation. Correlation coefficient and p-value were calculated using the Pearson correlation method, with p-values adjusted for multiple comparisons using the Benjamini & Hochberg method. **e,** Scatterplots of enriched TF motifs in the respective clusters. The y-axis shows the enrichment p-values of TF motifs, and the x-axis shows the average gene expression of the corresponding TFs.

To directly assess the impact of cell-type-specific methylation on TF binding, we performed motif enrichment analysis on differentially hypomethylated VMRs and examined the gene expression of each TF gene. Our findings uncovered a relationship between the TFs’ expression and the DNA methylation at their binding sites. For example, TFs associated with heart development, e.g., *Hand2*, *Tbx20*, and *Meis1*, were expressed in the cluster W6 whose VMRs are enriched in these TFs’ binding motifs (Fig. 2e (right)). Similar sets of tissue-specific TFs are found identified to other embryo structures, e.g., *Ebf1* and *Pbx1* in the brain/spinal cord and *Sox9*, *Ebf1*, and *Zeb2* in the craniofacial region (Fig. 2e). Of note, *Ebf1* was expressed, and its binding motif was also enriched across all three tissue regions (Fig. 2e). EBF1 was reported as an interaction partner for the TET2 and was suggested as a sequence-specific mechanism for regulating DNA methylation in cancer^35^.

To precisely resolve the fine structure of transcriptional regulation at the craniofacial and forebrain regions of the E11 embryo, we employed the 10 μm pixel-size microfluidic chip to produce a near single-cell resolution spatial map (Fig. 1g, h and Extended Data Fig. 10a). Integration of our spatial dataset with a single-cell RNA sequencing (scRNA-seq) reference^36^ from mouse embryo revealed a strong concordance between the two datasets (Extended Data Fig. 10b, c). For example, W11 maps to neuroectoderm and glial cells in the single-cell reference, W7 to CNS neurons, W5 to olfactory sensory neurons, and W10 to epithelial cells (Extended Data Fig. 10b, c). Cell-type deconvolution using the scRNA-seq reference^36^ identified distinct clusters that correspond precisely to known anatomical structures of the developing mouse brain. Notably, two spatially defined clusters, W7 and W11, captured key telencephalic compartments. W11 was enriched for telencephalon progenitors in the ventricular zone of the pallium, a neurogenic niche characterized by active cell division and proliferation, while W7 corresponded to GABAergic cortical interneurons localized in the mantle zone, where newborn neurons migrate, accumulate, and differentiate to establish cortical architecture^37-39^ (Fig. 1h and Extended Data Fig. 10d). Gene ontology (GO) enrichment analysis further supported these regional identities, highlighting biological processes associated with neurogenesis and progenitor proliferation in W11, and neuron projection and migration in W7 (Extended Data Fig. 10e). Moreover, the cell-type lineage tree constructed from the scRNA-seq reference confirmed the developmental trajectory, positioning telencephalon progenitors as direct precursors to GABAergic cortical interneurons^36^. Beyond the forebrain, our spatial analysis also resolved refined sensory structures within the developing olfactory system (cluster W5 and W10) (Fig. 1g and Extended Data Fig. 10a). Sensory neurons were notably enriched in W5 (Extended Data Fig. 10f, g), spatially localized adjacent to the forebrain. This spatial pattern closely aligns with the established developmental trajectory of the olfactory system, wherein olfactory sensory neurons progressively form connections with the forebrain as embryogenesis progresses^40-42^. Together, these findings provide a high-resolution view of neural and sensory system formation, underscoring the power of Spatial-DMT in resolving intricate anatomical structures and capturing the spatiotemporal dynamics.

### Dynamics of DNA methylation and gene expression during embryogenesis

Mammalian embryogenesis is a precisely timed process with dynamic DNA methylation and gene expression underlying cell differentiation and tissue development^26^, e.g., neuron tube formation^43^ and heart development^44^. By pseudo-time analysis and applying Spatial-DMT to embryos of two different gestational ages (E11 and E13), we can investigate the dynamics of DNA methylation, gene expression, and their interactions in both spatial and temporal scales. We first analyzed the developing brain, focusing on the differentiation trajectory from oligodendrocyte progenitors to premature oligodendrocytes. Spatial mapping of each pixel’s pseudo-time revealed an organized migration of oligodendrocyte progenitor cells from the subpallium to the pallium during oligodendrogenesis (Fig. 3a and Extended Data Fig. 9b). This pseudo-temporal process is associated with coupled DNA methylation and gene expression in different patterns. For example, DNA methylation loss can be both associated with gene activation (e.g., *Nrg3, an* oligodendrocyte marker^45^) and silencing (e.g., *Pdgfra*, an oligodendrocyte precursors markers^46^) (Fig. 3b, red arrows). The presence of both the positive and the negative couplings between DNA methylation and gene expression is aligned with the above comparisons across spatial pixels, reinforcing the regulatory diversity of mouse embryogenesis and oligodendrogenesis^46^.

**Fig. 3:**
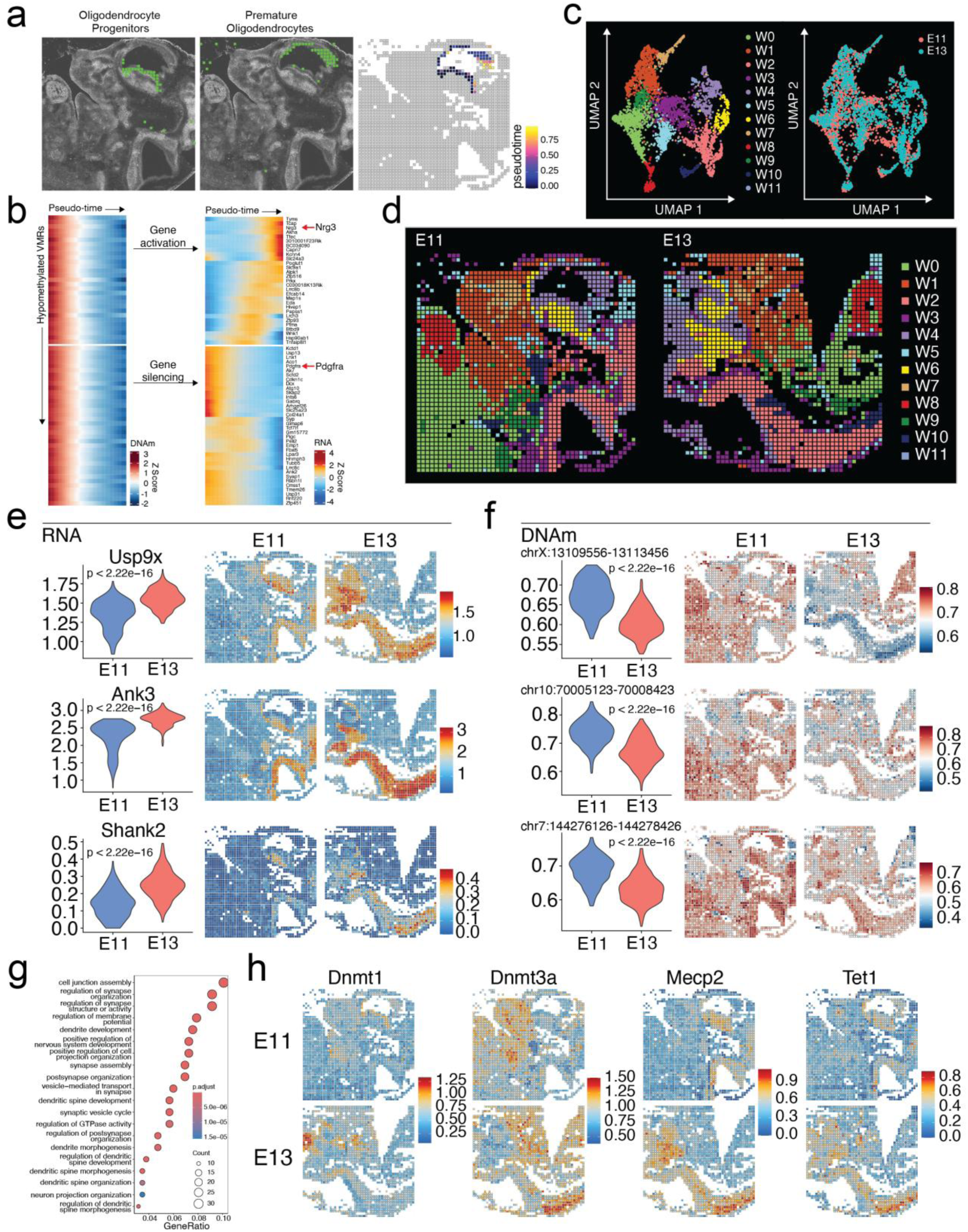
Spatiotemporal dynamics of DNA methylation and RNA transcription during embryogenesis. **a,** Spatial mapping of oligodendrocyte progenitors and premature oligodendrocytes identified by label transfer from mouse embryo scRNA-seq data^27^ to Spatial-DMT, with pseudotemporal reconstruction of the oligodendrogenesis plotted in space (right). **b,** Pseudo-time heatmaps of VMRs that become demethylated (left) and expression changes of nearby genes from oligodendrocyte progenitors to premature oligodendrocytes (right). Each row represents a specific genomic locus (left) and nearby genes (right), with columns representing tissue pixels ordered by pseudo-time. The color scale indicates the Z-scores of DNA methylation and gene expression. **c, d,** UMAP visualization **(c)** and spatial distribution **(d)** of integrated E11 and E13 WNN analysis. Spatial tissue pixels from different developmental stages conform well and match with the tissue types. **e, f,** Comparative analysis of RNA expression levels (log normalized expression) **(e)** and DNA methylation (methylation percentage) **(f)** for upregulated genes in E13 brain and spinal cord. **g,** GO enrichment analysis of biological processes related to demethylated and upregulated genes in the brain and spinal cord from E11 to E13 stages. **h,** Spatial mapping of the expression levels (log normalized expression) of DNA methylation-related enzymes in the brain and spinal cord regions across E11 and E13 stages.

To further illustrate the temporal molecular dynamics of embryonic development, we performed spatial mapping of the E13 mouse embryo (Extended Data Fig. 11a). First, we validated our RNA dataset by integrating it with a published spatial ATAC-RNA^18^ reference from E13 mouse embryo, which revealed a strong concordance (Extended Data Fig. 12a). Co-clustering the two datasets identified 11 distinct cluster populations, each corresponding to the histological location in both sample (Extended Data Fig. 12b, c). By integrating spatial data from E11 and E13 embryos (Fig. 3c, d), we identified genes upregulated in E13 (Fig. 3e and Extended Data Fig. 11b, d (left)) with notable DNA methylation loss (Fig. 3f, and Extended Data Fig. 11c, d (right)). These genes are implicated in the corresponding tissue functions. For example, *Usp9x*, *Ank3*, and *Shank2* are critically involved in neuronal development, synaptic organization, morphogenesis, and transmission^47-49^ (Fig. 3e, f). *Ctnna1*, *Pecam1*, and *Lamb1* are pivotal for maintaining cardiac tissue integrity^50^, vascular development^51^, and cardiac tissue structuring^52^, respectively (Extended Data Fig. 11b, c). Functional analysis of upregulated genes further corroborated their link to related biological processes, e.g., including synapse, neuron projection organization, and dendritic spine morphogenesis in the brain (Fig. 3g), and sphingolipid metabolic process in the heart (Extended Data Fig. 11e). Our tissue map data more precisely timed these gene and pathway activations to a specific stage of embryo development, tissue location and shed light on their epigenetic regulatory mechanisms. Intriguingly, besides regulators of tissue development, some DNA methylation writers^53^, readers^54^, and eraser enzymes^55^, e.g., *Dnmt1*, *Dnmt3a*, *Mecp2*, and *Tet1* in E13 embryo, showed higher expression in E13 embryo, suggesting elevated biochemical activity that drove the global DNA methylation dynamics (Fig. 3h).

### Spatial co-mapping of mCH, mCG, and transcription on postnatal mouse brain

Non-CpG cytosines methylation (mCH, where H denotes A, C, or T), particularly mCA, is uniquely abundant in the brain^22,25^. To evaluate the spatial heterogeneity of non-CpG cytosine methylation, we applied Spatial-DMT to the cortical and hippocampal regions of a postnatal 21-day-old (P21) mouse brain (Fig. 4a). Initial analysis of global mCA and mCG levels revealed relatively low methylation in the dentate gyrus (DG), cornu ammonis (CA)1/2, and CA3 regions, compared to the cortex (Extended Data Fig. 4d).

**Fig. 4:**
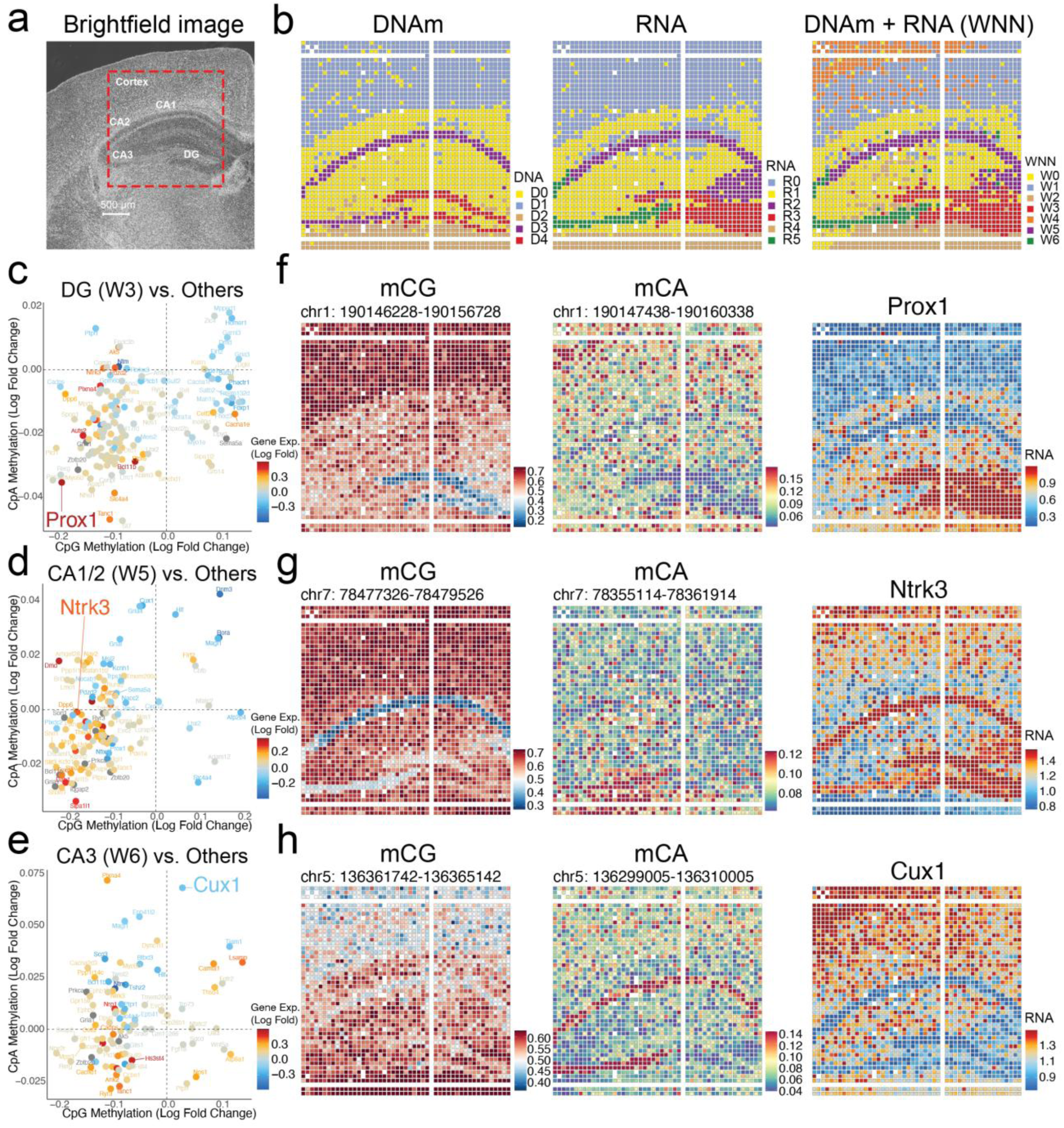
Spatial DNA methylation and RNA transcription analysis in the P21 mouse brain. **a,** Brightfield image of the P21 mouse brain section (pixel size, 20 μm, ROI area 2x2 mm^2^) showing the regions analyzed, including the DG, CA1, CA2, CA3, and cortex. **b,** Spatial distribution of clusters identified from DNA methylation data (left), RNA transcription data (middle), and integrated DNA and RNA data using WNN analysis (right). **c-e,** Scatter plot displaying the relationship between CpG and CpA methylation changes (log fold) and gene expression changes (log fold) for marker genes in DG **(c)**, CA1/2 **(d)**, CA3 **(e)**. The x-axis represents the log fold change in CpG methylation, and the y-axis represents the log fold change in CpA methylation. Each dot represents a specific gene, colored by the log fold change in gene expression, with red indicating upregulation and blue indicating downregulation. These plots demonstrate how differential methylation at CpG and CpA sites correlates with changes in gene expression, illustrating the complex regulatory landscape across different brain regions. **f-h,** Spatial mapping of CpG (left) and CpA (middle) methylation, and RNA expression levels (right) for selected genes in different brain regions.

The spatial distribution of methylome and transcriptome cluster matched anatomical structures shown by the histology image, reflecting the arealization of this brain region (Fig. 4b and Extended Data Fig. 13a). Clustering DNA methylation data yielded 5 clusters (D0-D4) (Fig. 4b (left) and Extended Data Fig. 13a (left) – DNAm). Similarly, RNA analysis revealed 6 clusters (R0-R5) (Fig. 4b (middle) and Extended Data Fig. 13a (middle) – RNA). Integrating both modalities refined the clustering (W0-W6) by additionally identifying the superficial cortical layer (W4) that was not detected by any single modality (Fig. 4b (right) and Extended Data Fig. 13a (right) – WNN). Known cell-specific marker genes are spatially distributed to regions that enrich these cell types. For instance, *Prox1*, a TF crucial for neurogenesis and the maintenance of granule cell identity^56^, was prominently expressed in DG (Fig. 4f (right)). *Satb1*, which plays a role in cortical neuron differentiation and layer formation^57^, was strongly enriched in the cerebral cortex (Extended Data Fig. 13b (right)). *Bcl11b*, a TF essential for neuronal progenitor cell differentiation^58^, demonstrated elevated expression in the hippocampus (Extended Data Fig. 13c (right)).

To elucidate the regulatory roles of mCG and mCA on gene transcription, we compared the two modifications on each cluster’s signature genes against all other clusters combined. By correlating the modification levels with gene expression, we identified genes potentially regulated by mCG, mCA, or both (Fig. 4c-e). *Prox1* and *Bcl11b* expression were significantly associated with both mCG and mCA (Fig. 4c, f, and Extended Data Fig. 13c), broadly marking the DG and CA1/2 regions, respectively. In contrast, *Ntrk3*, a receptor tyrosine kinase crucial for nervous system function^59^, was highly expressed in the hippocampal CA1/2 and DG regions, correlating primarily with mCG levels but not with mCA (Fig. 4d, g). Similarly, *Satb1* expression in the cortex was strongly correlated with mCG but not with mCA levels (Extended Data Fig. 13b). Lastly, the silencing of *Cux1*, a TF involved in neuronal development and function^60^, exhibited a negative correlation with CA and CG hypermethylation in the CA3 region. In contrast, in the CA1/2 region, *Cux1* expression showed a negative correlation only with CA hypermethylation and appeared independent of mCG levels (Fig. 4e, h). Across both sequence contexts, negative correlations between DNA methylation and gene expression are more prevalent than positive ones, highlighting the predominantly repressive nature of these epigenetic modifications (Extended Data Fig. 13d). Collectively, mCG and mCA regulate transcription in a gene-specific manner, jointly defining the cell identity across cell types and brain regions.

Further analysis of neuronal and glial populations revealed cell-type- and region-specific transcriptomic and epigenetic variation. For example, *Syt1* and *Rbfox3* are broadly expressed across neurons in all cortical layers (Extended Data Fig. 14a). In contrast, *Cux2*, *Cux1*, and *Satb2* were highly expressed in upper-layer neurons, whereas *Bcl11b* was enriched in the deeper cortical layers (Extended Data Fig. 4c). Oligodendrocytes (*Mbp* and *Plp1*) and fibrous astrocytes (*Gfap*) were specifically enriched in the corpus callosum and hippocampal regions (Extended Data Fig. 14b, d). To further resolve cell identities, we integrated our Spatial-DMT dataset with a reference scRNA-seq dataset^61^ to associate spatial clusters with previously defined cell types. For instance, W0, W3, and W5 were identified as oligodendrocytes, dentate gyrus granule neurons, and telencephalon-projecting excitatory neurons, respectively (Extended Data Fig. 15a-c). Aggregated gene expression and DNA methylation levels across pixels within each cluster were highly correlated with the corresponding single-cell RNA-seq transcriptomes^61^ (Extended Data Fig. 15d) and single-cell DNA methylome profiles^22^ (Extended Data Fig. 15e), as illustrated by W3 (DG) and W5 (CA1) (Fig. 4b (right)). Cell-type deconvolution using the same scRNA-seq reference^61^ revealed well-defined spatial organization of diverse cell types, consistent with known brain anatomy (Extended Data Fig. 15f). For example, cortical excitatory neurons showed expected laminar distribution: TEGLU7, TEGLU8, TEGLU4, and TEGLU3 were enriched in layers 2/3, 4, 5, and 6, respectively. In the hippocampus, TEGLU24 and TEGLU23 mapped to CA1/2 and CA3 regions, while granule neurons DGGRC2 localized to the dentate gyrus. Additionally, DEGLU1 was enriched in the thalamic region. All cell types were mapped to expected brain anatomical regions in agreement with the scRNA-seq reference^61^. These observations underscore the robustness of Spatial-DMT in resolving complex tissue architectures and simultaneously profiling epigenome and transcriptome with their spatial context, which remains challenging for conventional anatomical and single-cell approaches.

### Spatial-DMT resolves mitotic history and subtle region-specific epigenetic variations

Partially methylated domains (PMDs) are lamina-associated, late-replicating genomic regions that progressively lose methylation over mitotic divisions^62^, making them valuable indicators of a cell’s mitotic history. By spatially mapping PMD methylation in E11 and E13 embryos (Extended Data Fig. 16a, b), we identified distinct tissue-specific patterns. For instance, regions such as the forebrain and hindbrain/spinal cord exhibited higher PMD methylation, while embryonic heart tissue demonstrated lower PMD methylation levels, reflecting active cardiogenesis at these developmental stages^63^ (Extended Data Fig. 16a, b). Intriguingly, we observed spatial gradients in PMD methylation, decreasing from the center to the periphery in the heart, and from the mantle to the ventricular zone in the forebrain and hindbrain/spinal cord. These gradients potentially reflect the spatial organization of progenitor cells and their differentiation trajectories. In the postnatal P21 brain (Extended Figure 16c), cortical layers displayed higher PMD methylation, consistent with reduced proliferative capacity typical of differentiated neurons^64^. In contrast, the dentate gyrus exhibits comparatively lower PMD methylation, consistent with the presence of neural stem/progenitor cells in the subgranular zone that continues to undergo mitotic division and neurogenesis^65^.

The spatially resolved methylation landscape enables the identification of subtle, region-specific epigenetic variations that would otherwise be undetected. To illustrate this, we performed regional differential methylation analysis, comparing the ventricular and mantle zone of hindbrain/spinal cord (Extended Data Fig. 16d). This analysis identified differential methylation at neural progenitor cell-specific promoters and enhancers, as shown by the enrichment of corresponding H3K4me1 marks. These differential methylated loci colocalized with enhancers functionally implicated in brain and neural tube development (mEnhA9), which are associated with the transcription factors binding sites crucial for neuronal differentiation, including FOXO4^66^, NEUROG2^67^, and HOXC9^68^.

Spatially resolved methylation profiling further delineated epigenetic distinctions between brain subregions within a shared WNN cell cluster (Extended Data Fig. 16e). In the forebrain, regions exhibiting loss of methylation were enriched for binding sites of FOXI1, transcription factors crucial for auditory development^69^. In contrast, spinal cord-specific hypomethylation correlated with TLX1 occupancy, a transcription factor essential for spinal cord maturation and neuronal differentiation^70^. These findings demonstrate that Spatial-DMT can resolve subtle epigenetic heterogeneity across distinct tissue microenvironments.

Finally, the spatially resolved variations in methylome and transcriptome can complement each other in distinguishing differentiated cell states (Fig. 1f (left and middle)). Despite originating from the same RNA-defined clusters (R3), D0 and D4 cells exhibited distinct subpopulations when stratified by their VMRs. Motif enrichment analysis of low-methylation sites in D0 and D4 identified regulatory elements associated with facial and cardiac morphogenesis (Extended Data Fig. 16f), whereas corresponding gene expression changes were limited (Extended Data Fig. 16g). This may reflect epigenetically primed subpopulations that share similar transcriptional states. Together, these findings highlight the power of Spatial-DMT in delineating cell identity beyond the resolution of transcriptomic analysis alone.

## Discussion

Recent advances in single-cell DNA methylation and transcriptomic profiling have illuminated the diverse epigenetic architectures underlying cellular functions in both health and disease^1,22,26,71,72^. However, the anatomical context for DNA methylation and its regulatory effects on spatial cell organization within complex tissues remains to be elucidated.

In this study, we developed Spatial-DMT, a pioneering technology that enables simultaneous profiling of DNA methylation and RNA expression from cells within their native tissue context. The method yielded robust and high-quality data, demonstrating high reproducibility across technical replicates and strong concordance with established single-cell and spatial transcriptomic references. Using this reliable platform, we mapped the methylome of mouse embryos at E11 and E13, revealing developmental changes during embryogenesis. Extending our analysis to the postnatal brain, we profiled the mCH-enriched neuronal epigenome, consistent with previous findings of elevated mCA levels in mature neurons. By refining spatial resolution to 10 μm, we achieved near single-cell resolution, enabling the precise delineation of fine tissue architecture and the identification of region-specific epigenetic features. These regulatory elements, often difficult to resolve with traditional single-cell methods due to tissue dissociation, emerge clearly when the spatial context is preserved. Notably, spatial DNA methylation profiling provided unique information about cell mitotic history, a potential biomarker for cellular aging and oncogenesis. Importantly, the multi-modal nature of Spatial-DMT uncovered intricate relationships between DNA methylation and transcription within the tissue context. Integrating DNA methylation and transcription from the same tissue section delineated the cell-type and region-specific mechanisms of mCG and mCH regulation and their potential roles in transcriptional programs. Our data indicate that gene expression may be differentially influenced by mCG and mCA for different cell types in a spatially restricted manner, potentially due to the distinct genomic distribution of these methylation marks. While the observed associations are correlative, they offer valuable insights into the complex interplay of regulatory elements on gene expression. This integrative data analysis also demonstrated that each modality captures distinct yet complementary aspects of cellular states, enhancing cell states differentiation beyond what is achievable with single-modality approaches. This may reflect regulatory redundancy, where distinct epigenetic mechanisms converge to produce similar transcriptional outcomes. Conversely, the opposite scenario is equally plausible—transcriptional states may diverge despite similar epigenetic landscapes, as epigenetic regulation represents only one layer influencing gene expression. As a result, Spatial-DMT may reduce the need for high sequencing depth, offering a cost-effective strategy for unbiased genome-wide methylome analysis.

Future development areas include the following: (1) Our co-profiling approach lays the foundation that can be improved by incorporating additional omics layers with DNA methylation and RNA transcriptomes, improving our ability to unravel the complexities of gene regulation during development and disease progression. Potential integrations include chromatin conformation (HiC), chromatin accessibility (ATAC-seq), histone modifications (CUT&Tag), metabolome (mass spectrometry imaging), and surface proteins (CITE-seq). (2) Spatial-DMT, which is based on an Enzymatic Methyl-seq, does not distinguish between 5-methylcytosine and 5-hydroxymethylcytosine modification, the latter of which accumulates in certain brain regions^22^. This lack of resolution between distinct cytosine modifications complicates the interpretation of their respective roles in regulating gene expression. This limitation could be overcome by employing emerging methods that enable simultaneous measurement of the full spectrum of cytosine base modifications^73,74^. (3) Combining Spatial-DMT with long-read sequencing technologies, such as Oxford Nanopore Technologies and PacBio HiFi sequencing, can offer more direct, simplified, and comprehensive coverage of epigenetic modifications. (4) Given the stability of methylated DNA in formalin-fixed paraffin-embedded (FFPE) samples, extending our protocol for FFPE tissues could be valuable for clinical studies, with potential applications in precision medicine across diverse diseases. (5) Developing a user-friendly platform for spatial multi-omics analysis will streamline the analysis process, facilitating the exploration of DNA methylome, RNA transcriptome, and other omics data. (6) Lower-resolution pixels (e.g., 50 μm) generally capture more CpGs, UMIs, and expressed genes, likely due to the inclusion of a greater number of cells within each pixel. This effect depends on factors such as tissue heterogeneity and cell type variability and results in each pixel representing a mixture of signals from neighboring cells. Further development of advanced spatial computational deconvolution methods is essential for inferring single-cell resolution, minimizing potential biases, and enhancing biological interpretability. (7) Further optimization of the Spatial-DMT protocol may improve coverage and minimize potential tagmentation bias. (8) While this study primarily focuses on the mouse embryos and brain tissue, further validation across diverse tissues, developmental stages, and species will expand the applicability of this technology, which we identified as important future directions. Our current work lays a strong foundation for extending Spatial-DMT to broader biological contexts.

In summary, Spatial-DMT expands the spatial omics technologies by enabling the study of DNA methylation profiles and their interaction with the transcriptomes on the same tissue sections. Our datasets underscore the importance of the spatial tissue context when studying transcriptional regulation. We anticipate that this new spatial multi-omics technology will facilitate novel methylation biomarker discoveries, deepen our understanding of disease mechanisms, and benefit basic research and clinical applications.

## Methods

### Tissue slides preparation

Mouse C57 embryo sagittal frozen sections (no. MF-104-11-C57, MF-104-13-C57) were purchased from Zyagen. Freshly harvested E11 or E13 mouse embryos were snap-frozen in optimal cutting temperature compounds and sectioned at 7-10 μm thickness. Tissue sections were collected on poly-L-lysine-coated glass slides (Electron Microscopy Sciences, no. 63478-AS).

Juvenile mouse brain tissue (P21) was obtained from the C57BL/6 mice housed in the University of Pennsylvania Animal Care Facilities under pathogens-free conditions. All procedures used were pre-approved by the Institutional Animal Care and Use Committee.

Mice were sacrificed at P21 by CO_2_, followed by transcranial perfusion with cold DPBS. After isolation, brains were embedded in Tissue-Tek O.C.T. compound and snap-freezing on dry ice and 2-methylbutane bath. Coronal cryosections of 8-10 μm were mounted on the back of the Superfrost®/Plus microscope slides (Fisher Scientific, no. 12-550-15).

### Preparation of transposome

Unloaded Tn5 transposome (no. C01070010) were purchased from Diagenode, and the transposome was assembled following the manufacturer’s guidelines. The oligos applied for transposome assembly were: Tn5ME-B, 5’-/5Phos/CAT CGG CGT ACG ACT AGA TGT GTA TAA GAG ACA G-3’ Tn5MErev, 5’-/5Phos/CTG TCT CTT ATA CAC ATC T-3’

### DNA barcode sequences, DNA oligos, and other key reagents

DNA oligos used for PCR and library construction are shown in Supplementary Table 1. All DNA barcode sequences are provided in Supplementary Tables 2 (barcode A) and 3 (barcode B) and all other chemicals and reagents are in Supplementary Table 4.

### Fabrication of PDMS microfluidic device

Chrome photomasks were purchased from Front Range Photomasks, with a channel width of either 20 or 50 μm. The molds for polydimethylsiloxane (PDMS) microfluidic devices were fabricated using standard photolithography. The manufacturer’s guidelines were followed to spin-coat SU-8-negative photoresist (Microchem, nos. SU-2025 and SU-2010) onto a silicon wafer (WaferPro, no. C04004). The heights of the features were about 20 and 50 μm for 20- and 50-μm -wide devices, respectively. PDMS microfluidic devices were fabricated using the SU-8 molds. We mixed the curing and base agents in a 1:10 ratio and poured the mixture onto the molds. After degassing for 30 min, the mixture was cured at 66-70 °C for 2-16 h. Solidified PDMS was extracted from the molds for further use. The detailed protocol for the fabrication and preparation of the PDMS device can be found in our publication ^24^.

### Spatial joint profiling of DNA Methylation and RNA Transcription

Frozen tissue slides were quickly thawed for 1 min in a 37 °C incubator. The tissue was fixed with 1% formaldehyde in PBS containing 0.05 U ml ^-1^ RNase inhibitor (Enzymatics) for 10 min and quenched with 1.25 M glycine for another 5 min at room temperature. After fixation, tissue was washed twice with 1 ml of DPBS-RI and cleaned with deionized (DI) H_2_O.

Tissue was then permeabilized with 100 μl of 0.5 % triton X-100 plus 0.05 U ml ^-1^ RNase inhibitor (RI) for 30 min at room temperature, then washed twice with 200 μl DPBS-RI for 5 min each. After permeabilization, tissue was treated with 100 μl of 0.1 N HCl for 5 min at room temperature to disrupt histones from the chromatin, then washed twice with 200 μl of wash buffer (10 mM Tris-HCl pH7.4, 10 mM NaCl, 3 mM MgCl_2_, 1% BSA, 0.1% Tween 20) plus 0.05 U ml ^-1^ RNase inhibitor for 5 min at room temperature. 50 μl of transposition mixture (5 μl of assembled transposome, 16.5 μl of 1x DPBS, 25 μl of 2x Tagmentation buffer, 0.5 μl of 1% digitonin, 0.5 μl of 10% Tween-20, 0.05 U ml^-1^ RNase inhibitor (Enzymatics), 1.87 μl of nuclease-free H_2_O) was added and incubated at 37°C for 60 min. After 60 min incubation, the first round of transposition mixture was removed, and a second round of 50 μl of fresh transposition mixture was added and incubated for another 60 min at 37°C. To stop the transposition, 200 μl of 40 mM EDTA with 0.05 U ml^-1^ RNase inhibitor was added with incubation for 5 min at room temperature. After that 200 μl 1x NEB3.1 buffer plus 1% RNase inhibitor was used to wash the tissue for 5 min at room temperature. The tissue was then washed again with 200 μl of DPBS-RI for 5 min at room temperature before proceeding with the In-situ Reverse Transcription (RT) reaction.

### In-situ Reverse Transcription (RT)

For the In-situ RT, the following mixture was added: 12.5 μl 5x RT buffer, 4.05 μl RNase-free water, 0.4 μl RNase inhibitor (Enzymatics), 1.25 μl 50% PEG-8000, 3.1 μl 10mM dNTPs, 6.2 μl 200 U μl^-1^ Maxima H Minus Reverse Transcriptase, 25 μl 0.5x DPBS-RI, and 10 μl 100μM RT primer (Biotinylated-poly-deoxythymidine (dT) oligo). The tissue was incubated for 30 min at room temperature, then at 45°C for 90 min in a humidified container. After the RT reaction, tissue was washed with 1x NEB3.1 buffer plus 1% RNase inhibitor for 5 min at room temperature.

### In-situ barcoding

For the first barcode (barcode A) in situ ligation, the first PDMS chip was covered to the tissue region of interest. For alignment purposes, a 10x objective lens (KEYENCE BZ-X800 fluorescence microscope, BZ-X800 Viewer Software) was used to take the brightfield image. The PDMS device and tissue slide were clamped tightly with a homemade acrylic clamp. Barcode A was first annealed with ligation linker 1; mix 10 μl of 100 μM ligation linker, 10 μl of 100 μM individual barcode A, and 20 μl of 2x annealing buffer (20 mM Tris-HCl pH7.5-8.0, 100 mM NaCl_2_, 2 mM EDTA). For each channel, 5 μl of ligation master mixture was prepared with 4 μl of ligation mixture (27 μl T4 DNA ligase buffer, 0.9 μl RNase inhibitor (Enzymatics), 5.4 μl 5% Triton X-100, 11 μl T4 DNA ligase, 71.43 μl RNase free water), 1 μl of each annealed DNA barcode A (A1-A50, 25 μM). The vacuum was applied to flow the ligation master mixture into 50 channels of the device and cover the region of interest of the tissue, followed by incubation at 37°C for 30 min in a humidified container. Then the PDMS chip and clamp were removed after washing the tissue with 1x NEB 3.1 buffer for 5 min. The slide was then washed with deionized water and dried with compressed air.

For the second barcode (barcode B) In-situ ligation, the second PDMS chip was covered to the region of interest and a brightfield image was taken with the 10x objective lens. An acrylic clamp was applied to clamp the PDMS and tissue slide together. Annealing of barcode B (B1-B50, 25 μM) and preparation of the ligation mixture are the same as barcode A. The whole device was incubated at 37°C for 30 min in a humidified container. The PDMS chip and clamp were then removed, and the slide was washed with deionized water and dried with compressed air. A brightfield image was then taken for further alignment.

### Reverse crosslinking

For tissue lysis, the region of interest was digested with 100 μl of the reverse crosslinking mixture (0.4 mg ml^-1^ proteinase K, 1 mM EDTA, 50 mM Tris-HCl pH8.0, 200 mM NaCl, 1% SDS) at 58-60°C for 2 h in a humidified container. The lysate was then collected in a 0.2 ml PCR tube and incubated on a 60°C shaker overnight.

### gDNA and cDNA separation

For gDNA and cDNA separation, the lysate was purified with Zymo DNA Clean & Concentrator kit and eluted with 100 μl nuclease-free water. 40 μl of Dynabeads MyOne Streptavidin C1 beads were used and washed 3 times with 1x B&W buffer containing 0.05% Tween-20 (50 μl 1M Tris-HCl pH8.0, 2000 μl 5M NaCl, 10 μl 0.5 M EDTA, 50 μl 10% Tween-20, 7890 μl nuclease-free water). After washing, beads were resuspended in 100 μl of 2x B&W buffer (50 μl 1M Tris-HCl pH8.0, 2000 μl 5M NaCl, 10 μl 0.5 M EDTA, 2940 μl nuclease-free water) containing 2 μl of SUPERase In RNase inhibitor, then mixed with the gDNA/cDNA lysate and allowed binding for 1 h with agitation at room temperature. A magnet was then used to separate the beads, which bind the cDNA that contains biotinylated deoxythymidine (dT), from the supernatant that contains the gDNA.

### gDNA library generation

200 μl of supernatant was collected from the above separation process for further methylated gDNA detection and library construction. 1 ml of DNA binding buffer was added to the 200 μl supernatant and purified with the Zymo DNA Clean & Concentrator kit again, then eluted in 84 μl (3x 28 μl) of nuclease-free water. The NEBNext^®^ Enzymatic Methyl-seq conversion module (EM-seq^TM^) was then used to detect methylated DNA in the sample by converting unmethylated cytosines to uracil, the manufacturer’s guidelines were followed. 28 μl of DNA sample was aliquoted to each PCR tube, TET2 reaction mixture (10 μl TET2 Reaction Buffer containing reconstituted TET2 Reaction Buffer Supplement, 1 μl Oxidation Supplement, 1 μl DTT, 1 μl Oxidation Enhancer, 4 μl TET2) was added to the DNA sample on ice. 5 μl of diluted 1:1300 of 500 mM Fe (II) solution was added to the mixture and incubated for 1 h at 37°C in a thermocycler. After the reaction, transfer the sample to ice add 1 μl of Stop Reagent from the kit, and incubate for another 30 min at 37°C. TET2 converted DNA was then purified with 90 μl of SPRI beads and eluted with 16 μl nuclease-free water. Pre-heat the thermocycler to 85°C, mix 4 μl formamide to the converted DNA, and incubate for 10 min at 85°C in the pre-heated thermocycler. After the reaction, immediately place the heated sample on ice to maintain the open chromatin structure, then add the following reagents from the kit (68 μl nuclease-free water, 10 μl APOBEC Reaction Buffer, 1 μl BSA, 1 μl APOBEC) to deaminate unmethylated cytosines to uracil for 3 h at 37°C in a thermocycler. Deaminated DNA was then cleaned up using 100 μl (1:1 ratio) of SPRI beads and eluted in 20 μl of nuclease-free water.

### Splint ligation

The gDNA tube was heat-shocked for 3 min at 95°C and put on ice for 2 min immediately. Then add 10 μl of 0.75 μM pre-annealed Splint Ligate P5 (SLP5) adapter (diluted from 12 μM stock, which contained mix of 6 μl 100 μM SLP5RC oligo, 8.4 μl 100 μM SLS5ME-A-H10 oligo, 5 μl 10x T4 RNA Ligase Buffer, 30.6 μl nuclease-free water in PCR tube, and incubated 95°C for 1 min, then cool down gradually -0.1°C/sec to 10°C on thermocycler), 80 μl of ligation master mixture (40 μl preheated 50% PEG 8000, 12.5 μl SCR buffer (666 mM Tris-HCl pH8.0, 132mM MgCl2 in nuclease-free water), 10 μl 100 mM DTT, 10 μl 10 mM ATP, 1.25 μl 10,000 U/ml T4 PNK, 6.25 μl 400,000 U ml^-1^ T4 ligase) was added to the gDNA tube at room temperature. The splint ligation mixture was then splinted into 5 x 0.2 ml PCR tubes, 20 μl/tube. The tubes were shaken at 1000 rpm for 10s and spun down, then incubated for 45 min at 37°C, followed by 20 min at 65°C to inactivate the ligase. For splint ligation indexing PCR, 80 μl of the PCR reaction mixture (20 μl 5x VeraSeq GC Buffer, 4 μl 10 mM dNTPs, 3 μl VeraSeq Ultra Enzyme, 5 μl 20x EvaGreen dye, 2 μl 10 μM N501 primer, 2 μl 10 μM N70X-HT primer (Supplementary Table 1)) was mixed to each splint ligated tube. The mixture was then aliquoted to a new PCR tube with 50 μl volume and run on a thermocycler with the setting below, 98°C for 1 min, then cycling at 98°C for 10 s, 57°C for 20 s, and 72°C for 30 s, 13-19 cycles, followed by 72°C for 10 s. The reaction was removed once the qPCR signal began to plateau. The amplified PCR products were pooled and purified with 0.8X volume ratio of SPRI beads, and the completed DNA library was eluted in 15 μl of nuclease-free water.

### cDNA library generation

The separated beads contained cDNA were used for cDNA library generation. 400 μl of 1x B&W buffer with 0.05% Tween-20 was used to wash the beads twice, then the beads were washed once with 400 μl 10 mM Tris-HCl pH8.0 containing 0.1% Tween-20 for 5 min at room temperature. Streptavidin beads with bound cDNA molecules were placed onto a magnetic rack and washed once with 250 μl of nuclease-free water before being resuspended in a TSO solution (44 μl 5x Maxima RT buffer, 44 μl of 20% Ficoll PM-400 solution, 22 μl of dNTPs, 5.5 μl of 100 mM template switch oligo, 11 μl Maxima H Minus Reverse Transcriptase, 5.5 μl of RNase inhibitor (Enzymatics), and 88 μl of nuclease-free water). Resuspended beads were then incubated for 30 min with agitation at room temperature, and then for 90 min at 42°C, with gentle agitation. After the reaction, beads were washed with 400 μl of 10 mM Tris pH8.0 containing 0.1% Tween-20, then washed without resuspension in 250 μl nuclease-free water. Remove water on the magnetic rack and resuspend the beads in the PCR solution (100 μl 2x Kappa Master mix, 8.8 μl 10 μM primer 1 and 2, 92.4 μl Nuclease-free water). Mix well and split 50 μl of the PCR mixture into 4 individual 0.2 ml PCR tubes. Run the PCR program: 95°C for 3 min and cycling at 98°C for 20 s, 65°C for 45 s, and 72°C for 3 min, total 5 cycles, followed by 4°C on hold. After 5 cycles of PCR reaction, place the 4x PCR tubes onto a magnetic rack and transfer 50 μl of the clear PCR solution to 4x independent optical grade qPCR tubes, adding 2.5 μl of 20X Evagreen dye to each tube, and run the sample on qPCR machine with the following conditions: 95°C for 3 min, cycling at 98°C for 20 s, 65°C for 20 s and 72°C for 3 min, 13-17 cycles, followed by 72°C for 5 min. The reaction was removed once the qPCR signal began to plateau. The amplified PCR product was purified with 0.8x volume ratio of SPRI bead and eluted in 20 μl of nuclease-free water.

A Nextera XT DNA Library Prep Kit was used for cDNA library preparation. 2 μl (2 ng) of purified cDNA (1 ng μl^-1^), 10 μl Tagment DNA buffer, 5 μl Amplicon Tagment mix, and 3 μl nuclease-free water was mixed and incubated at 55°C for 5 min. 5 μl of NT buffer was then added to stop the reaction with incubation at room temperature for 5 min. PCR master mix (15 μl 2x N.P.M. Master mix, 1 μl of 10 μM P5 primer (N501) and 1 μl of 10 μM indexed P7 primer (N70X), 8 μl of nuclease-free water) was added, and run the PCR reaction with the following program: 95°C for 30 s, cycling at 95°C for 10 s, 55°C for 30 s, 72°C for 30 s and 72°C for 5 min, total for 12 cycles. The PCR product was then purified with a 0.7X ratio of SPRI beads and eluted in 15 μl nuclease-free water to obtain the cDNA library.

### Library quality check and NGS sequencing

An Agilent Bioanalyzer D5000 ScreenTape was used to determine the size distribution and concentration of the library before sequencing. NGS was conducted on an Illumina NovaSeq 6000 sequencer and NovaSeq X Plus system (150bp paired-end mode).

### Data preprocessing

For RNA sequencing data, Read 2 was processed to extract barcode A, barcode B, and the Unique Molecular Identifiers (UMIs). Using the STARsolo pipeline (version 2.7.10b)^75^, this processed data was mapped to the mouse genome reference (mm10). This step generated a gene matrix that captures both gene expression and spatial positioning information, encoded through the combination of barcodes A and B. The gene matrix was then imported into R for downstream spatial transcriptomic analysis using Seurat package (version 5.1.0)^76^.

For DNA methylation data, adaptor sequences were trimmed before demultiplexing the fastq files using the combination of barcodes A and B. We employed the BISulfite-seq CUI Toolkit (BISCUIT) (version 0.3.14)^77^ to align the DNA sequences to the mouse reference genome (mm10). Methylation levels at individual CG and CH sites were stored as continuous values between 0 and 1, representing the fraction of methylated reads after quality filtering. These processed CG/CH files were then independently analyzed using the MethSCAn pipeline^28^ to identify variably methylated regions (VMRs), defined as fused genome intervals with methylation level variance in the top 2 percentile. We used default parameter settings when running MethSCAn, and the MethSCAn filter min-sites parameter was determined from the read coverage knee plot (Extended Data Fig. 4a). The methylation levels and residuals of VMRs were then imported into R for downstream DNA methylation analysis.

### Clustering and data visualization

We first identified the location of pixels on tissue from the brightfield image using a custom python script (https://github.com/zhou-lab/Spatial-DMT-2024/), before removing additional empty barcodes based on read count thresholds determined by the knee plot (Extended Data Fig. 4a).

For RNA data, we used the SCTtransform function in Seurat package (version 5.1.0), built upon a regularized negative binomial model, for normalization and variance stabilization. Dimensionality reduction was performed using RunPCA function with the SCTtransformed assay. We then constructed the nearest neighbor graph using the first 30 principal components (PCs) with the FindNeighbors function and identified clusters with the default Leiden method in the FindClusters function. Finally, a UMAP embedding was computed using the same PCs with RunUMAP function.

Due to the inherent sparsity of DNA methylation data, it is impractical to analyze methylation status solely at the individual CpG level. Binary information at sparse loci cannot be directly used to construct a feature matrix suitable for downstream analysis. In our study, we adopted the Variably Methylated Regions (VMRs) framework, which divides the genome into variable-sized tiles and calculates the average methylation level across CpGs within each tile for each pixel^78^. This approach results in a continuous-valued matrix, where rows correspond to pixels and columns represent genomic tiles, with values ranging from 0 to 1. VMR methylation levels and residuals were then imputed using the iterative PCA approach as suggested in the MethSCAn instructions. Initially, missing residual values were replaced with zero, and missing methylation levels were replaced with the average values for that VMR interval. The PCA approach was iteratively applied until updated values stabilized to a threshold. The imputed residual matrix for VMRs was then imported into the existing Seurat object as another modality. Similar to the RNA clustering pipeline, dimensionality was reduced using RunPCA function. The first 10 PCs from the residual matrix were used for clustering and UMAP embedding.

To visualize clusters in their spatial locations, the SpatialDimPlot function was used after clustering by gene expression or VMR residual. UMAP embedding was visualized with the DimPlot function. The FindMarkers function was applied to select genes and VMRs that were differentially expressed or methylated for each cluster. For spatial mapping of individual VMR methylation levels or gene expression, we applied the smoothScoresNN function from the FigR package^79^. The SpatialFeaturePlot function was then used to visualize VMR methylation level and gene expression across all pixels. To illustrate the relationships between clustering results from different modalities, we generated the confusion matrix and alluvial diagram using the pheatmap and ggalluvial R package^80^.

### Integrative data analysis

To integrate spatial DNA methylation and RNA data, Weighted Nearest Neighbors (WNN) analysis in Seurat was applied using the FindMultiModalNeighbors function. Based on the weighted nearest neighbors graph, clustering, UMAP embedding, and spatial mapping of identified clusters were performed for integrated visualization.

For the integration of mouse embryo E11 and E13 spatial transcriptomics data, the top 3000 integration features were selected, followed by using PrepSCTIntegration and IntegrateData functions to generate an integrated dataset. Similarly, to integrate with public single-cell transcriptomic data^36,61^, we first identified anchors using the FindIntegrationAnchors function in Seurat, followed by data integration using the IntegrateData function. To integrate DNA methylation data, common VMRs between both developmental stages were obtained, and the integrated CCA method from the IntegrateLayers function was used to join the methylation data from the two developmental stages. Wilcoxon signed-rank test was performed to compare the methylation levels and gene expression differences between the two timepoints.

### Transcription factor motif enrichment

To perform transcription factor motif enrichment, we first used the MethSCAn diff function on distinct groups of cells to identify differentially methylated VMRs based on the clustering assignment. The HOMER^81^ findMotifsGenome function was then applied to conduct known transcription factor motifs enrichment analysis using its default database. We followed the same parameter settings used in MethSCAn, with motif lengths of 5, 6, 7, 8, 9, 10, 11, and 12.

### CpGs enrichment analysis

Enrichment analysis of individual CpGs in the differential regions (Extended Figure 16d, e) was performed using knowYourCG (https://github.com/zhou-lab/knowYourCG), which provides a comprehensive annotation database for each CpG, including chromatin states, transcription factor binding sites, motif occurrences, PMD annotations, and more. To avoid inflated odds ratios for high coverage data, genomic uniformity was quantified using fold enrichment, defined as the ratio of observed overlaps to expected overlaps. The expected number of overlaps was calculated as: (number of CpGs sequenced × number of CpGs in the chromatin state feature) / total number of CpGs in the genome.

### Correlation and GO enrichment analysis

Correlation analysis was performed for different clusters. We first used the findOverlaps function in GenomicRanges package (version 4.4)^82^ to map VMRs to overlapped genes. Then, the Pearson correlation test was applied to obtain the correlation between mapped genes and corresponding VMRs. The Benjamini-Hochberg procedure was used to adjust all P-values.

GO enrichment analysis was conducted using the enrichGO function from clusterProfiler package (version 4.2)^83^. For GO enrichment in the comparative analysis of E11 and E13 mouse embryos, the FindMarkers function in Seurat package was used to find differential genes and VMRs in the same cluster from integrated data across two developmental stages. Differentially upregulated genes (FDR ≤ 0.05) with demethylated VMRs were used for GO analysis.

## Data availability

Raw and processed data reported in this paper are deposited in the Gene Expression Omnibus (GEO) with accession code GSE270498. Resulting fastq files were aligned to the mouse reference genome (mm10). Published data for data quality comparison and integrative data analysis include single cell atlas of mouse embryos (https://oncoscape.v3.sttrcancer.org/atlas.gs.washington.edu.mouse.rna/downloads, https://omg.gs.washington.edu/), mouse brain atlas (http://mousebrain.org/adolescent/downloads.html), and Allen Mouse Brain Atlas (https://developingmouse.brain-map.org/).

## Code availability

The data analysis pipeline and code to reproduce analyses are available on GitHub (https://github.com/zhou-lab/Spatial-DMT-2024/).

## Acknowledgements

We acknowledge the support from the Packard Fellowship for Science and Engineering (to Y.D.), the pilot award from the Epigenetics Institute at the University of Pennsylvania (to Y.D.), and the US National Institutes of Health (grant number DP2AI177913 to Y.D.), National Institute of Health / National Institute of General Medical Sciences [R35-GM146978] (to W.Z.). C.N.L. was supported in part by the Institute for RNA Innovation of the Perelman School of Medicine at the University of Pennsylvania.

## Contributions

Methodology: C.N.L. and Y.D.; Experimental Investigation: C.N.L. and A.C.; Data Analysis: C.N.L., H.F., W.Z., and Y.D.; Original Draft: C.N.L., H.F., W.Z., and Y.D. All authors reviewed, edited, and approved the manuscript.

---

**Supplementary Table 1.**
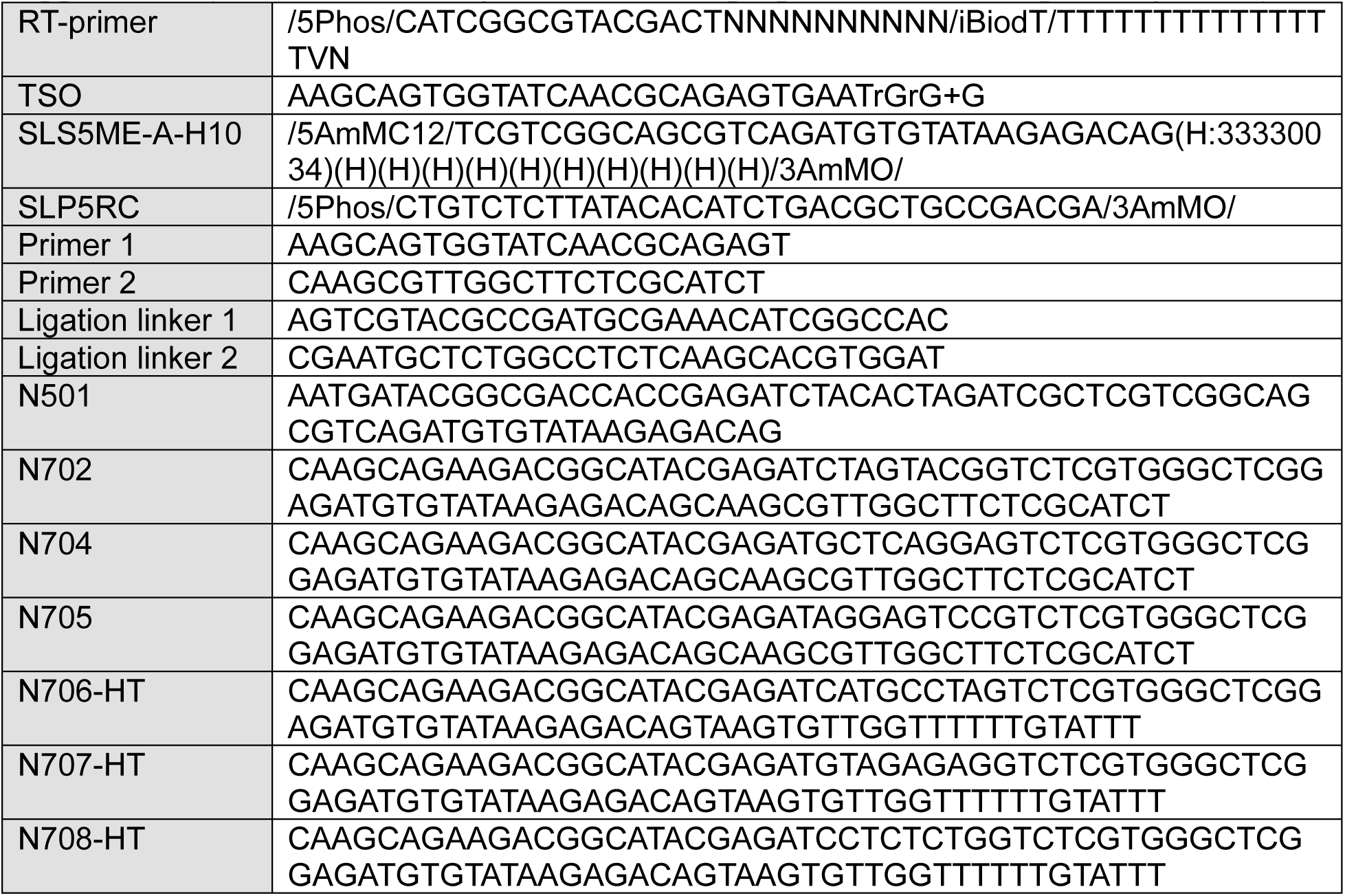
DNA oligos for PCR and preparation of the sequencing library.

**Supplementary Table 2.**
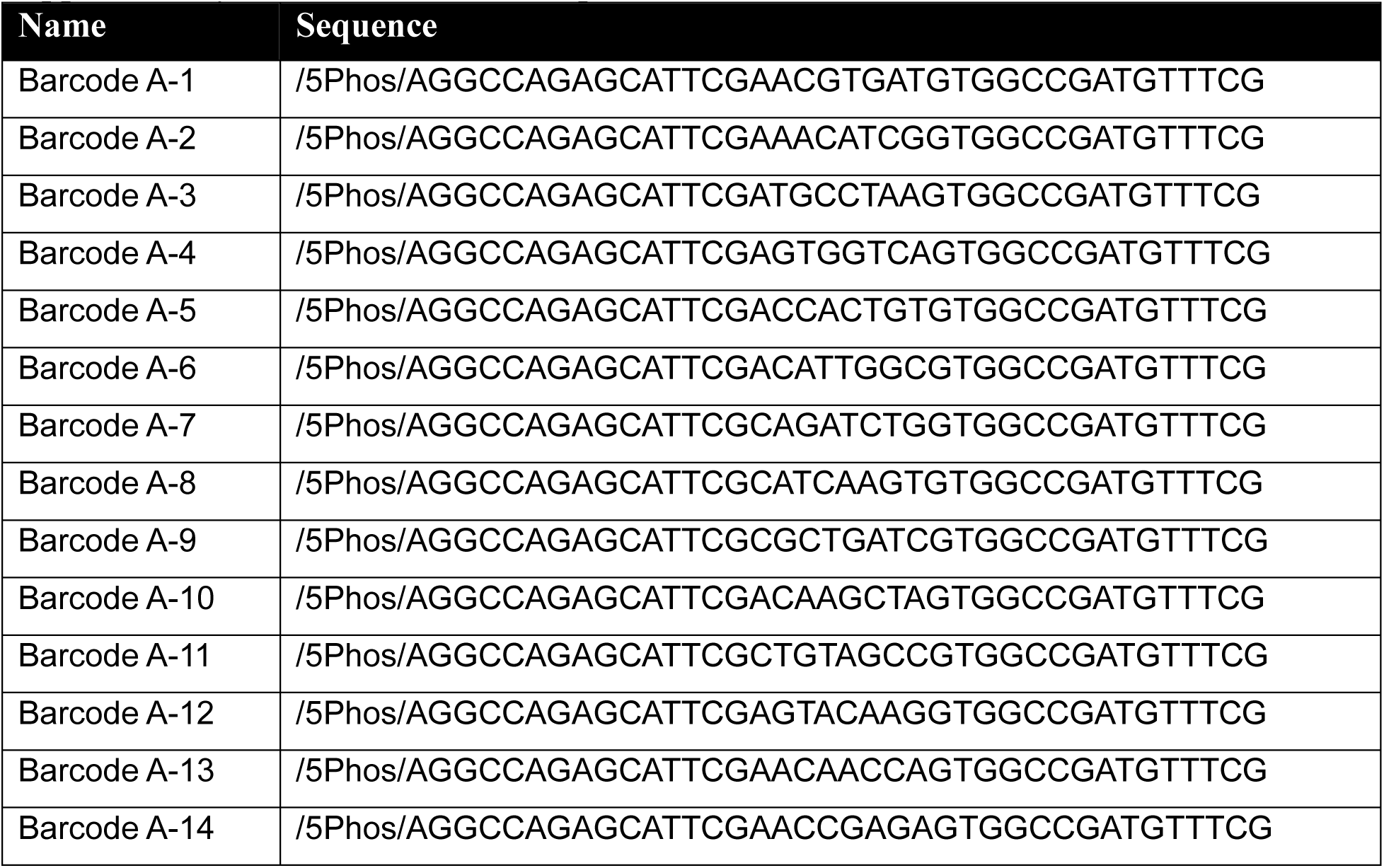
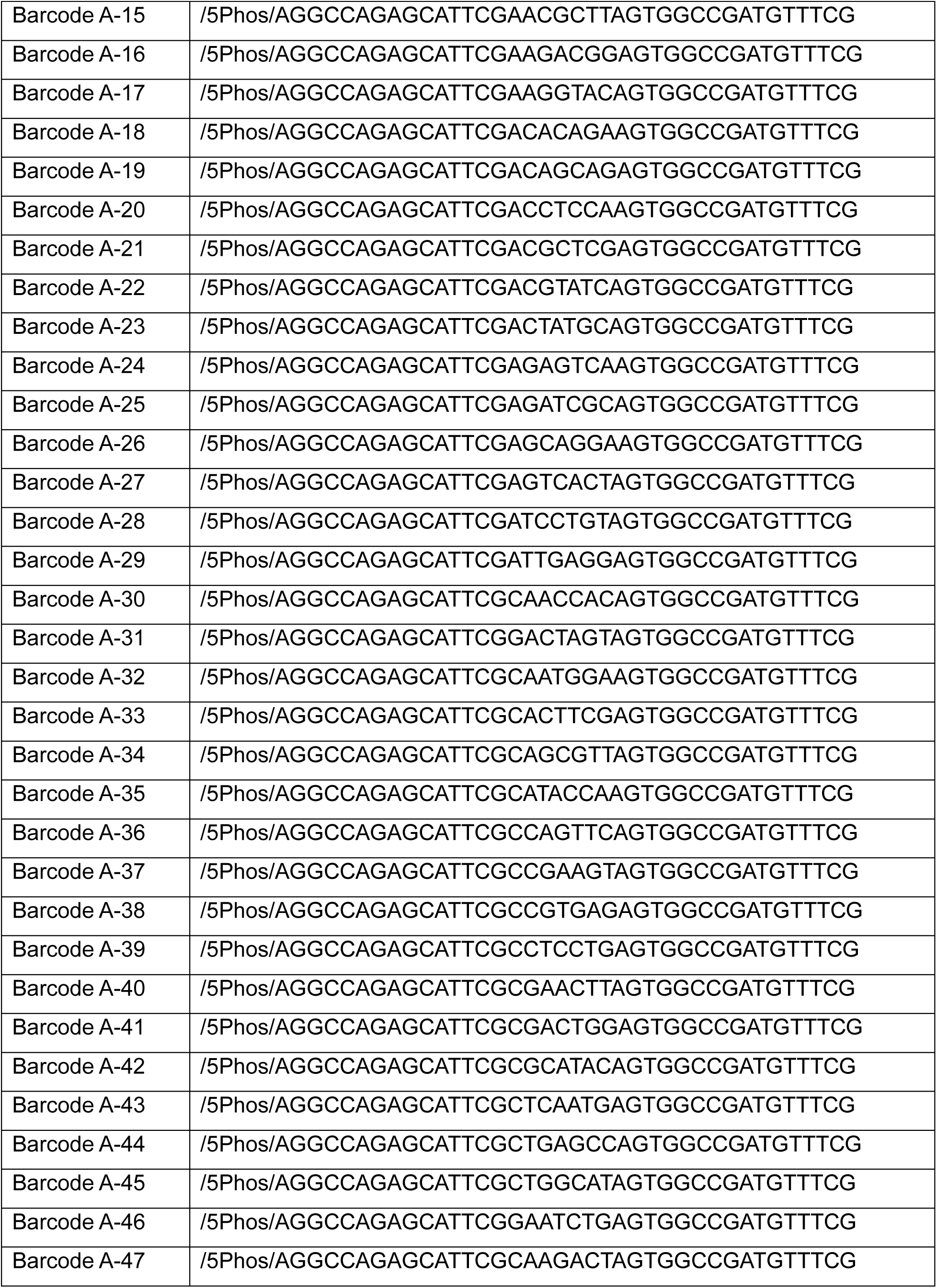
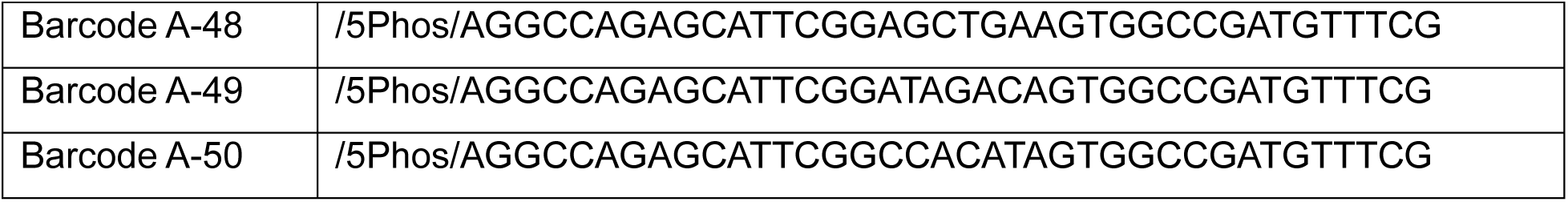
Barcode A Sequence.

**Supplementary Table 3.**
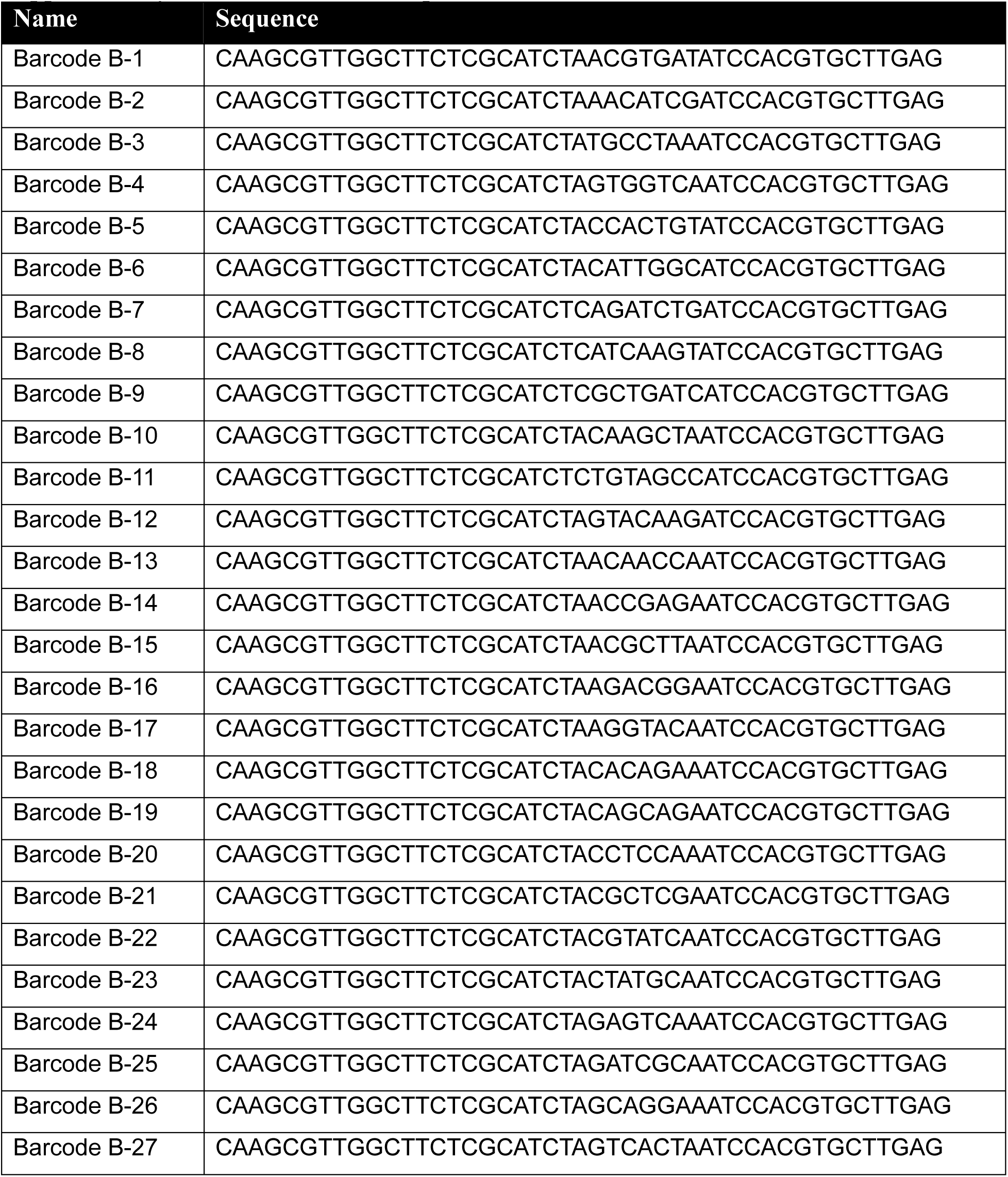
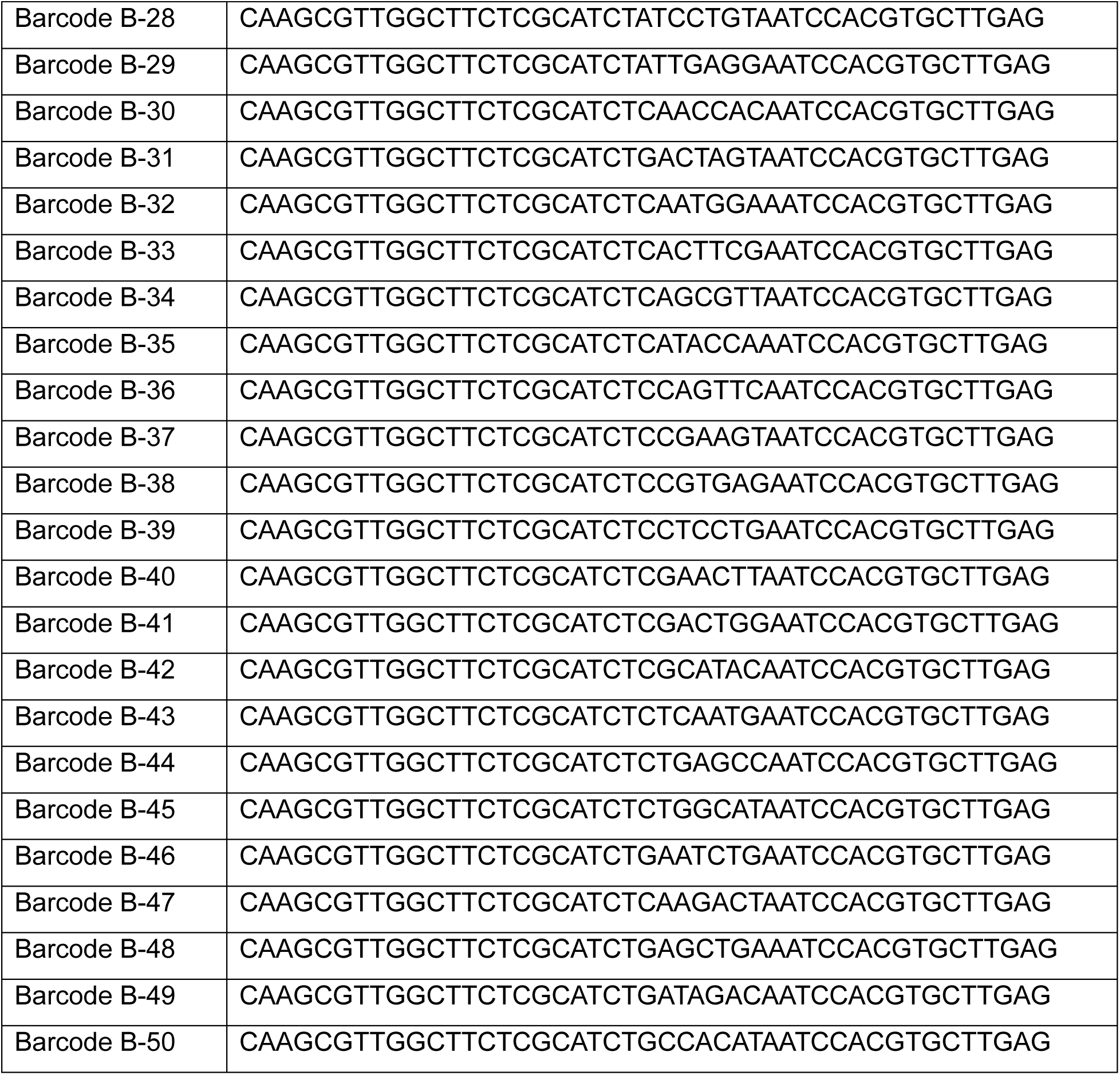
Barcode B Sequence.

**Supplementary Table 4.** Chemicals and reagents.

| Name | Catalog number | Vender |
| --- | --- | --- |
| Formaldehyde solution | PI28906 | Thermo Fisher Scientific |
| Glycine | 50046 | Sigma-Aldrich |
| 0.1N HCl | 2104-50ML | Sigma-Aldrich |
| NaCl | AM9760G | Thermo Fisher Scientific |
| MgCl<sub>2</sub> | AM9530G | Thermo Fisher Scientific |
| Digitonin | G9441 | Promega |
| Sodium dodecyl sulfate | 71736 | Sigma-Aldrich |
| HEPES pH 7.5 | BBH-75-250 | Boston BioProducts |
| EDTA Solution pH 8.0 | AB00502 | AmericanBio |
| Bovine Serum Albumin (BSA) | A8806 | Sigma-Aldrich |
| NP40 | 11332473001 | Sigma-Aldrich |
| Triton X-100 | T8787 | Sigma-Aldrich |
| T4 DNA Ligase | M0202L | New England Biolabs |
| T4 DNA Ligase Reaction Buffer | B0202S | New England Biolabs |
| NEBuffer 3.1 | B7203S | New England Biolabs |
| DPBS | 14190144 | Thermo Fisher Scientific |
| Proteinase K | EO0491 | Thermo Fisher Scientific |
| NEBNext High-Fidelity 2X PCR Master Mix | M0541L | New England Biolabs |
| SYBR Green I Nucleic Acid Gel Stain | S7563 | Thermo Fisher Scientific |
| DNA Clean & Concentrator-5 | D4014 | Zymo Research |
| Tn5 Transposase - unloaded | C01070010 | Diagenode |
| Tagmentation Buffer (2x) | C01019043 | Diagenode |
| Maxima H Minus Reverse Transcriptase (200 U/μl) | EP0751 | Thermo Fisher Scientific |
| VeraSeq Ultra DNA polymerase | 7520L | Qiagen |
| dNTP mix | R0192 | Thermo Fisher Scientific |
| SUPERase•In™ RNase Inhibitor (RI) | AM2694 | Thermo Fisher Scientific |
| RNase Inhibitor (40 U/μl) | Y9240L | Enzymatics |
| SPRI beads | A63880 | Beckman Coulter |
| Dynabeads MyOne C1 | 65001 | Thermo Fisher Scientific |
| Kapa Hotstart HiFi ReadyMix | KK2601 | Kapa Biosystems |
| NEBNext® Enzymatic Methyl-seq Conversion Module | E7125S | New England Biolabs |
| Nextera XT DNA Library Preparation Kit | FC-131-1024 | Illumina |

**Supplementary Table 5.** Summary of metrics for DNA methylation and RNA transcription co-profiling of all samples.

| Sample | E11 (10 μm) |  | E11#1 (50 μm) |  | E11#2 (50 μm) |  |
| --- | --- | --- | --- | --- | --- | --- |
| Modality | DNA | RNA | DNA | RNA | DNA | RNA |
| Total raw reads | 2,756,840,516 | 172,492,101 | 2,864,519,160 | 98,539,436 | 2,863,772,201 | 350,172,191 |
| Total pixels | 2493 | 2493 | 1954 | 1954 | 1947 | 1947 |
| Retained read % | 32.20 | 70.71 | 65.72 | 64.78 | 61.89 | 68.06 |
| Retained reads (before alignment) | 887,671,712 | 121,967,091 | 1,882,630,968 | 63,833,555 | 1,772,395,274 | 238,318,000 |
| Ave. retained reads / pixel | 355,069 | 12,476 | 753,052 | 7,544 | 708,958 | 37,810 |
| Total # of genes | - | 25,118 | - | 25,992 | - | 27,306 |
| Ave. gene # / pixel | - | 1,890 | - | 2,417 | - | 4,626 |
| Ave. UMI # / pixel | - | 3,596 | - | 5,108 | - | 16,709 |
| Ave. mCG % coverage | 0.62 | - | 1.09 | - | 1.29 | - |
| Ave. mCG # / pixel | 136,639 | - | 238,040 | - | 281,447 | - |
| Ave. mCH % coverage | - | - | - | - | - | - |
| Ave. mCH # / pixel | - | - | - | - | - | - |

| Sample | E13 (50 μm) | | P21 (20 μm) | |
| --- | --- | --- | --- | --- |
| Modality | DNA | RNA | DNA | RNA |
| Total raw reads | 3,018,600,847 | 180,508,321 | 3,914,623,812 | 110,320,779 |
| Total pixels | 1699 | 1699 | 2235 | 2235 |
| Retained read % | 55.69 | 72.65 | 42.48 | 73.88 |
| Retained reads (before alignment) | 1,680,936,562 | 131,147,184 | 1,662,805,699 | 81,501,569 |
| Ave. retained reads / pixel | 672,375 | 15,715 | 665,122 | 11,205 |
| Total # of genes | - | 28,695 | - | 23,822 |
| Ave. gene # / pixel | - | 3,788 | - | 2,525 |
| Ave. UMI # / pixel | - | 9,830 | - | 5,796 |
| Ave. mCG % coverage | 1.17 | - | 0.89 | - |
| Ave. mCG # / pixel | 255,402 | - | 194,224 | - |
| Ave. mCH % coverage | - | - | 0.43 | - |
| Ave. mCH # / pixel | - | - | 4,773,740 | - |

**Supplementary Table 6.** Summary of significant chromHMM full stack annotations.

| Name | Description |
| --- | --- |
| mEnhA2 | Active enhancers in most cell and tissue types (e.g. brain, heart, digestive, limb, epithelium) weaker in some cell types (e.g. liver, spleen, immune). Strong DNase, ATAC, H3K4me1, H3K27ac, H3K9ac. |
| mEnhA4 | Active enhancers signals (DNase, H3K4me1, H3K27ac, ATAC) in multiple tissue types including kidney, lung, heart, stomach, limb. |
| mEnhA5 | Weak to moderate enhancer signal (H3K4me1, H3K27ac, DNase, ATAC) in multiple tissues including heart, stomach, kidney, intestine, lung, and liver. |
| mEnhA7 | Strong H3K4me1 and DNase in brain, epithelium/neural tube, limb, and embryo. Lower H3K4me1 and DNase in others. |
| mEnhA8 | H3K4me1 and DNase in brain and epithelium/neural tube, also low H3K4me1 and DNase in various other tissues. |
| mEnhA9 | Enhancers in brain and epithelium/neural tube; H3K4me1, H3K27ac, DNase, ATAC in those tissue. |
| mEnhA10 | Strong enhancers in brain and epithelium/neural tube with H3K4me1, H3K27ac, H3K9ac, DNase, ATAC in those tissues. |
| mEnhA12 | Active enhancer signals (ATAC, DNase, H3K4me1, H3K27ac) in many tissue types. Weaker or less active enhancer signals in some tissue types including brain, epithelium/neural tube, connective tissue, and embryo. |
| mEnhA16 | Active enhancer mark (H3K27ac) strongest in limb, embryo. Moderate DNase in a subset of other cell types (limb, epithelium, musculature, ESC, gonad, adrenal gland, stomach, intestine, kidney, heart, brain). |
| mPromF4 | Promoter state enriched flanking TSS. Stronger H3K4me2 than H3K4me1 and H3K4me3. Low to moderate ATAC, DNase, H3K4me1/2/3, H3K9ac, |
| mPromF7 | Poised promoters/proximal enhancers particularly in brain and epithelium/neural tube; Strongest signals for H3K4me2 in Brain and Neural tubes and higher relative expression in those tissues; |
| openC1 | DNase is the strongest mark in state and found varying levels for different cell and tissue types. Some weak H3K27me3 and H3K4me1. |
| openC2 | Weak DNase in various cell types. Weaker H3K4me1 and H3K27me3 in a subset. |

**Extended Data Fig. 1:**
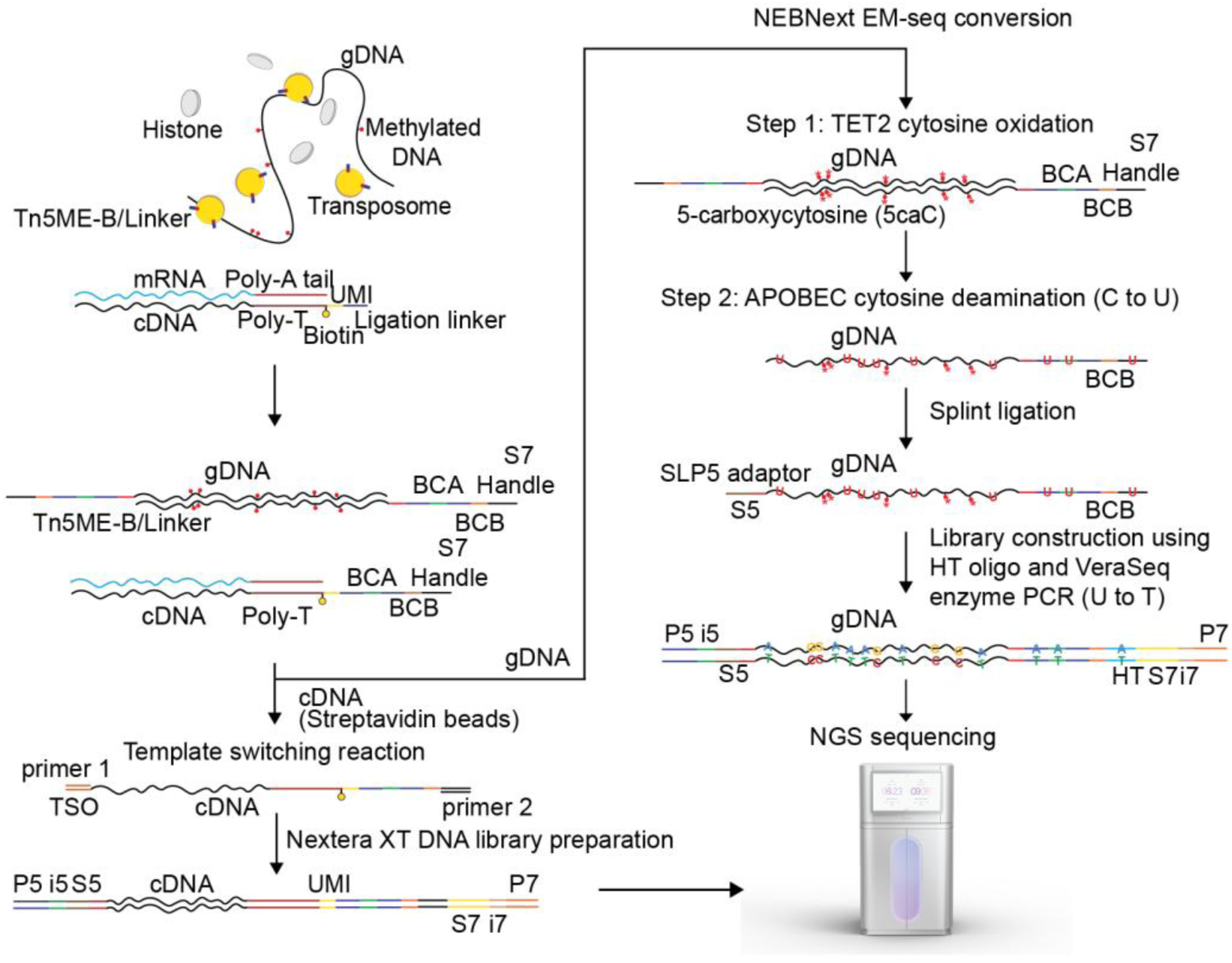
Chemistry workflow of Spatial-DMT. The diagram illustrates the chemistry workflow for simultaneous DNA methylation and RNA library preparation in Spatial-DMT. Initially, tissue sections are fixed and permeabilized to preserve tissue architecture and molecular integrity while allowing reagent penetration. Following permeabilization, sections undergo HCl treatment to disrupt the protein structure and remove nucleosome histones to improve Tn5 transposome accessibility. Next, Tn5 transposition integrates adapters into genomic DNA, and in situ reverse transcription (RT) converts mRNA into complementary DNA (cDNA) directly within the tissue. Subsequently, Barcode A and Barcode B are sequentially ligated to label and spatially encode DNA and mRNA molecules. Streptavidin bead separation then facilitates the differential enrichment and processing of nucleic acids: cDNA molecules undergo a template switching reaction, followed by Nextera XT library preparation, while genomic DNA (gDNA) is subjected to NEBNext EM-seq conversion for methylation detection, splint ligation, and subsequent library construction.

**Extended Data Fig. 2:**
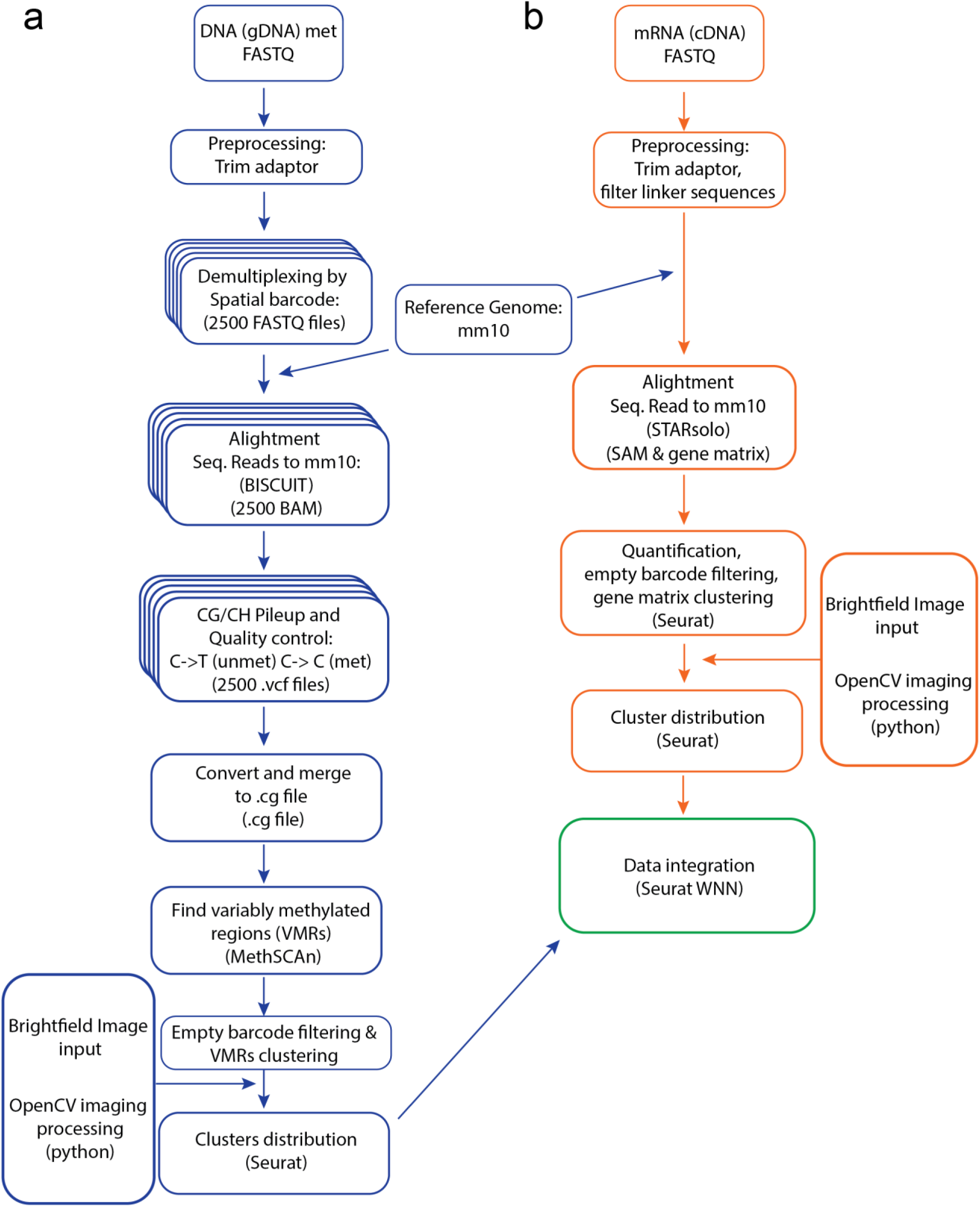
Computational workflow for Spatial-DMT data processing and analysis. The workflow diagram illustrates the steps involved in processing and analyzing spatial DNA methylation and RNA transcription data from Spatial-DMT. For DNA methylation, the process includes FASTQ preprocessing (trimming of adaptors and filtering of linker sequences), demultiplexing of spatial barcodes for 2500 FASTQ files, alignment of sequencing reads to the reference genome (e.g., mm10) using BISulfite-seq CUI Toolkit (BISCUIT) (resulting in 2500 BAM files), CG/CH pileup and quality control (variant calling to generate 2500 .vcf files and then conversion and merging to .cg files), identification of variably methylated regions (VMRs) using MethSCAn, empty barcode filtering and VMR clustering, and visualization of spatial clusters using Seurat, by integrating with brightfield images for spatial context processed by OpenCV. For mRNA data, the process includes FASTQ preprocessing (trimming of adaptors and filtering of linker sequences), alignment of sequencing reads to the reference genome (e.g., mm10) using STARsolo (generating SAM and gene matrix files), quantification of gene expression, filtering of empty barcodes, and visualization of spatial clusters using Seurat, by integrating with brightfield images for spatial context processed by OpenCV. The final step involves the integration of DNA methylation and RNA transcription data using Seurat WNN analysis to achieve integrated spatial multi-omics profiles.

**Extended Data Fig. 3:**
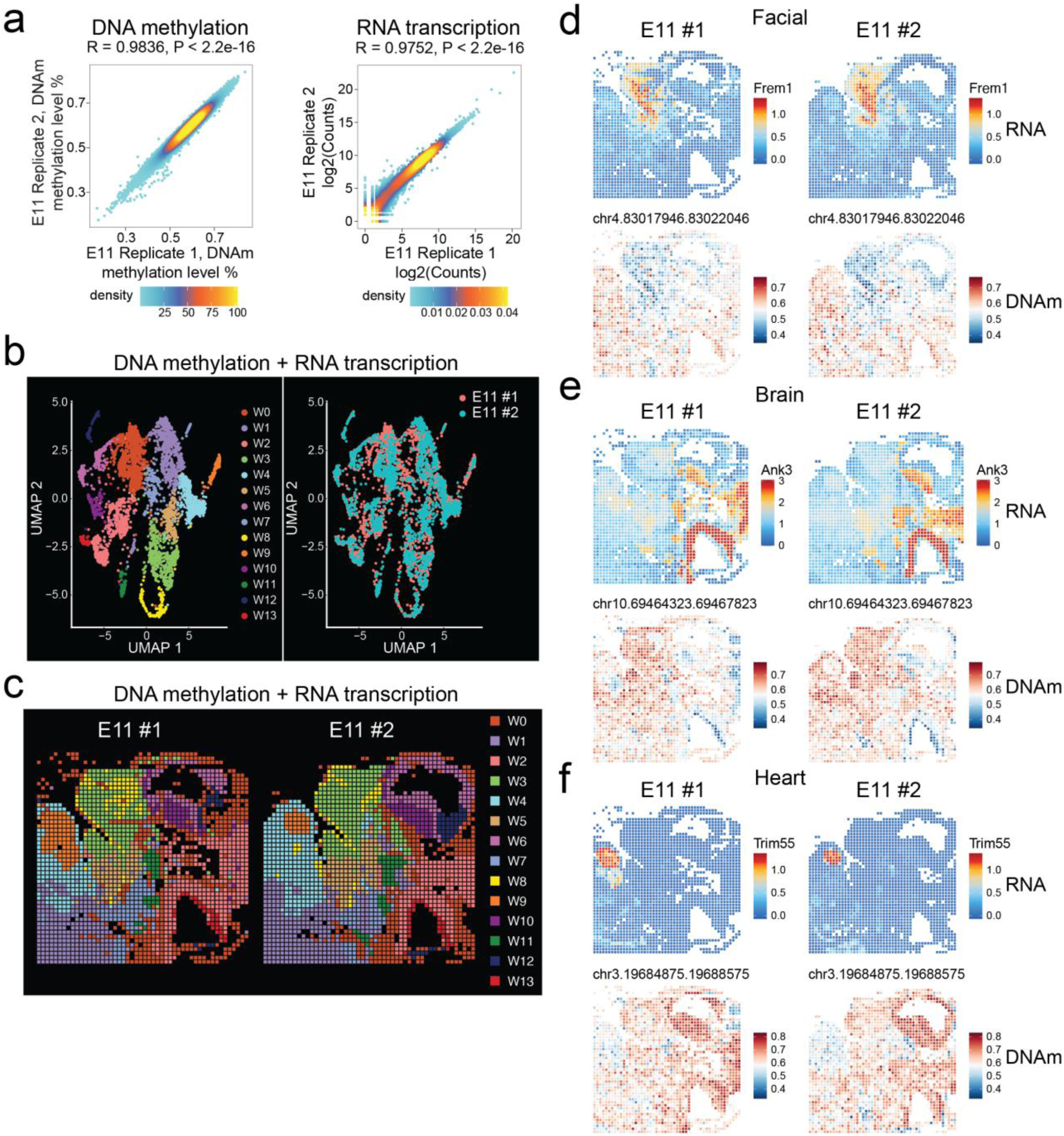
Reproducibility of Spatial-DMT. **a,** Correlation analysis between replicates based on DNA methylation (left) and RNA transcription (right) data from E11 embryo samples. **b, c,** Integrative analysis **(b)** and spatial distribution **(c)** of clusters from two independent Spatial-DMT experiments using E11 embryo samples. **d-f,** Spatial mapping of DNA methylation and RNA expression levels for selected marker genes in facial **(d)**, brain **(e)**, and heart **(f)** regions from two independent Spatial-DMT experiments using E11 embryo samples.

**Extended Data Fig. 4:**
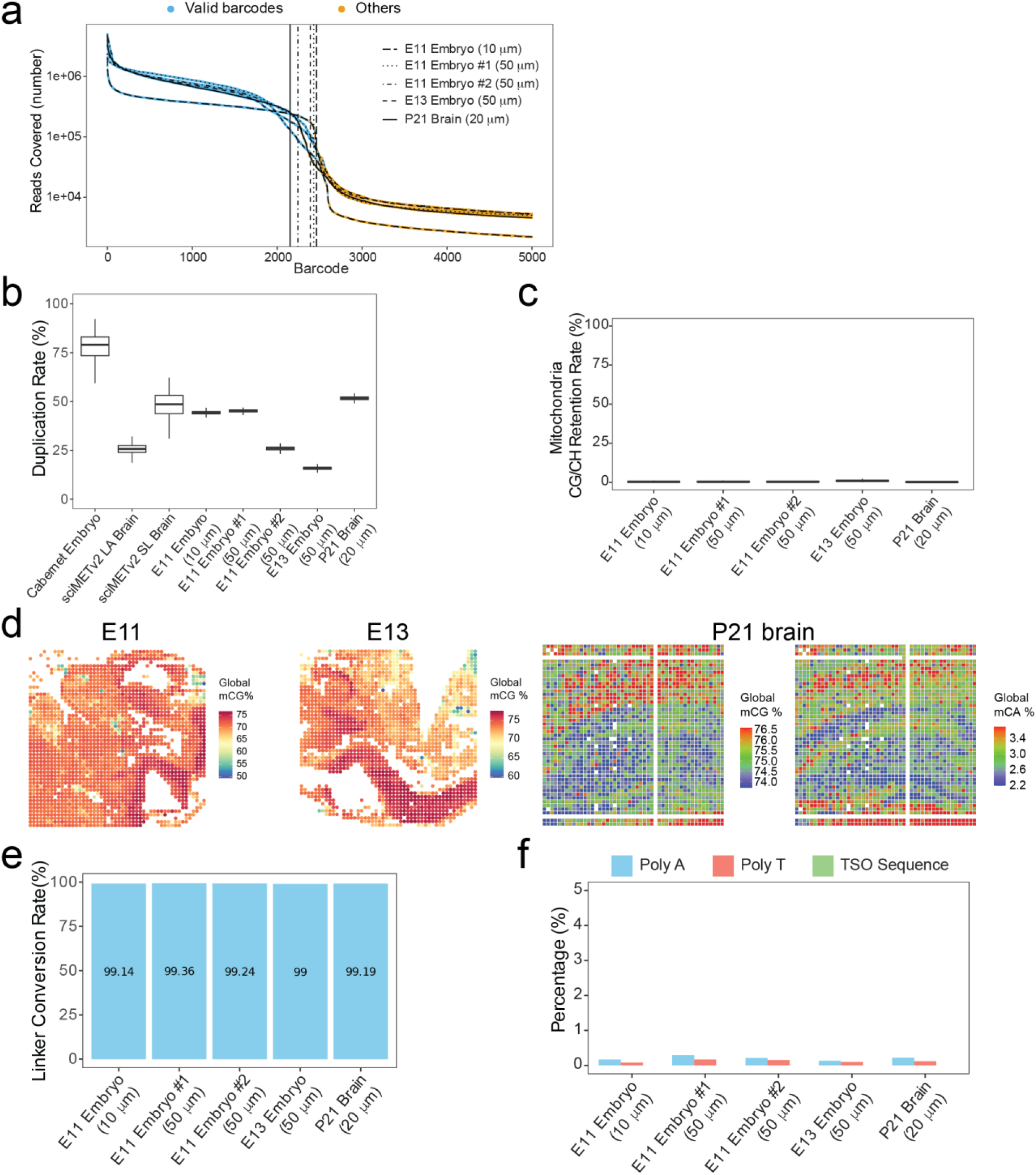
Quality control metrics for Spatial-DMT datasets. **a,** Barcode rank plots showing the distribution of reads per spatial barcode for E11, E13 embryos, and P21 brain samples. Valid barcodes are shown in blue, and filtered barcodes are shown in orange. **b,** Box plots showing the duplication rate for E11, E13 embryos, and P21 brain samples, and other single-cell DNA methylation datasets^16,85^. The y-axis represents the percentage of duplicated reads. **c,** Box plots showing the mitochondrial retention rate for E11, E13 embryos, and P21 brain samples. The y-axis represents the percentage of reads mapped to the mitochondrial genome. **d,** Spatial distribution heatmaps of global DNA methylation levels in E11, E13 embryos (mCG), and P21 mouse brain (mCG and mCA). **e,** The conversion rate of unmethylated cytosine in the linker sequence in E11, E13 embryos, and P21 mouse brain. **f,** Percentage of reads with poly A (≥ 30 adenine), poly T (≥ 30 thymine), and TSO sequence (AAGCAGTGGTATCAACGCAGAGTACATGGG) in the DNA libraries of E11, E13 embryos, and P21 mouse brain.

**Extended Data Fig. 5:**
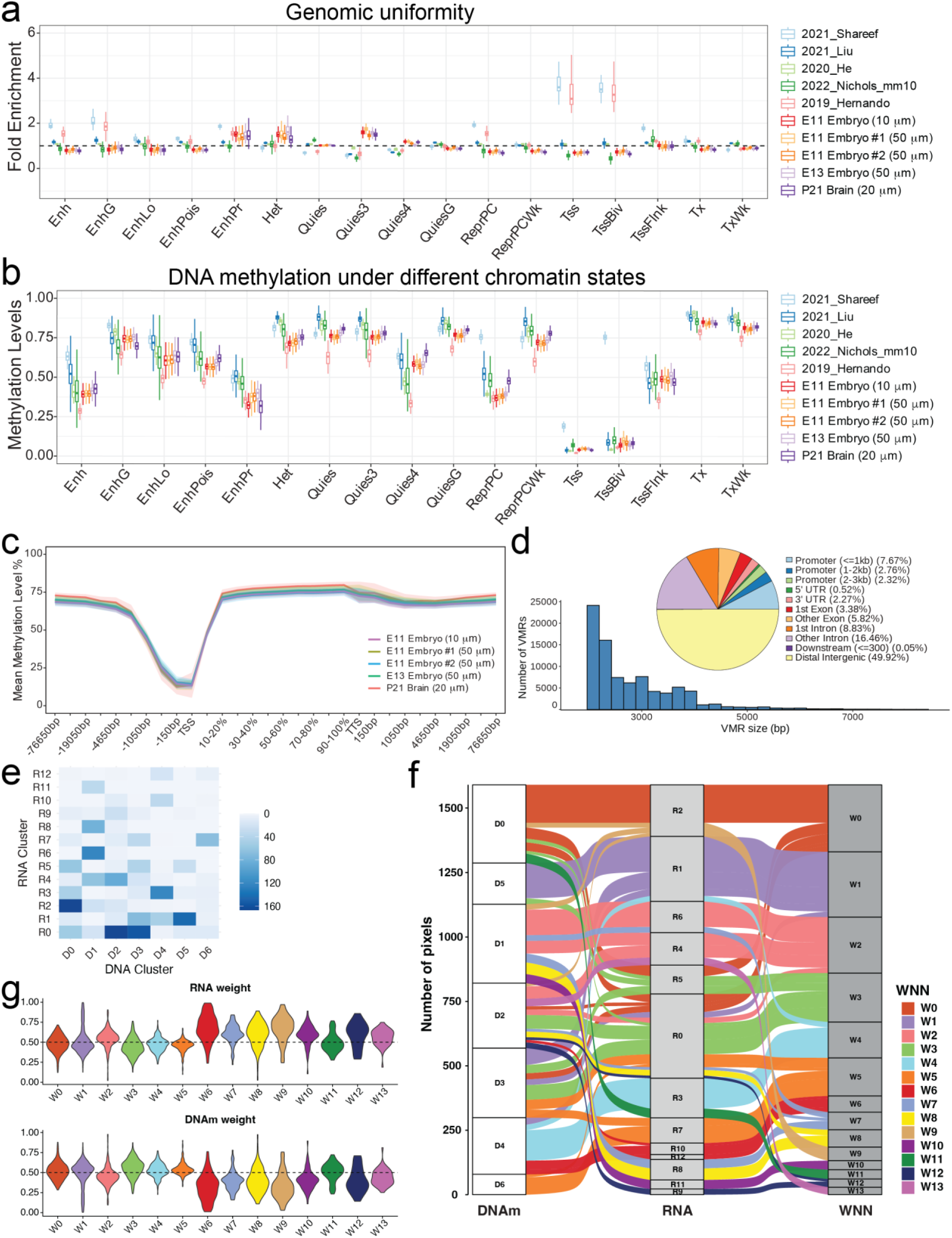
Further analysis of DNA methylation data quality of Spatial-DMT. **a,** Genomic uniformity analysis showing the fold enrichment (observed overlaps divided by expected overlaps) for different genomic features, comparing E11, E13 embryos, and P21 brain samples with reference datasets^7,16,22,23,26^. **b,** DNA methylation levels under different chromatin states, comparing E11, E13 embryos, and P21 brain samples with reference datasets^7,16,22,23,26^. The y-axis represents the methylation levels, with different chromatin states on the x-axis. These chromatin states include active states such as active transcription start site (TSS)-proximal promoter states (TssFlink), actively-transcribed states (Tx, TxWk), enhancer states (Enh, EnhG, EnhLo, EnhPois, EnhPr). Inactive states consist of constitutive heterochromatin (Het), quiescent states (Quies, Quies3, Quies4, QuiesG), repressed Polycomb states (ReprPC, ReprPCWk), and bivalent regulatory states (TssBiv). **c,** Line plots showing the mean DNA methylation levels (%) across different genomic regions for E11, E13 embryos, and P21 brain samples. The x-axis represents the genomic regions, and the y-axis represents the mean methylation levels. **d,** Histogram and pie chart showing the size distribution and genomic annotation of variably methylated regions (VMRs) identified in the E11 embryo. The pie chart indicates the percentage of VMRs in different genomic features, such as promoters, exons, and intergenic regions. **e,** Confusion matrix illustrating the correspondence between clustering results derived from RNA transcription and DNA methylation data in the E11 embryo. **f,** Alluvial diagram showing the relationships among clusters identified by DNA methylation, RNA, and WNN integration in the E11 embryo. **g,** Modality weights indicating the relative contributions of gene expression and DNA methylation for each WNN cluster in the E11 embryo.

**Extended Data Fig. 6:**
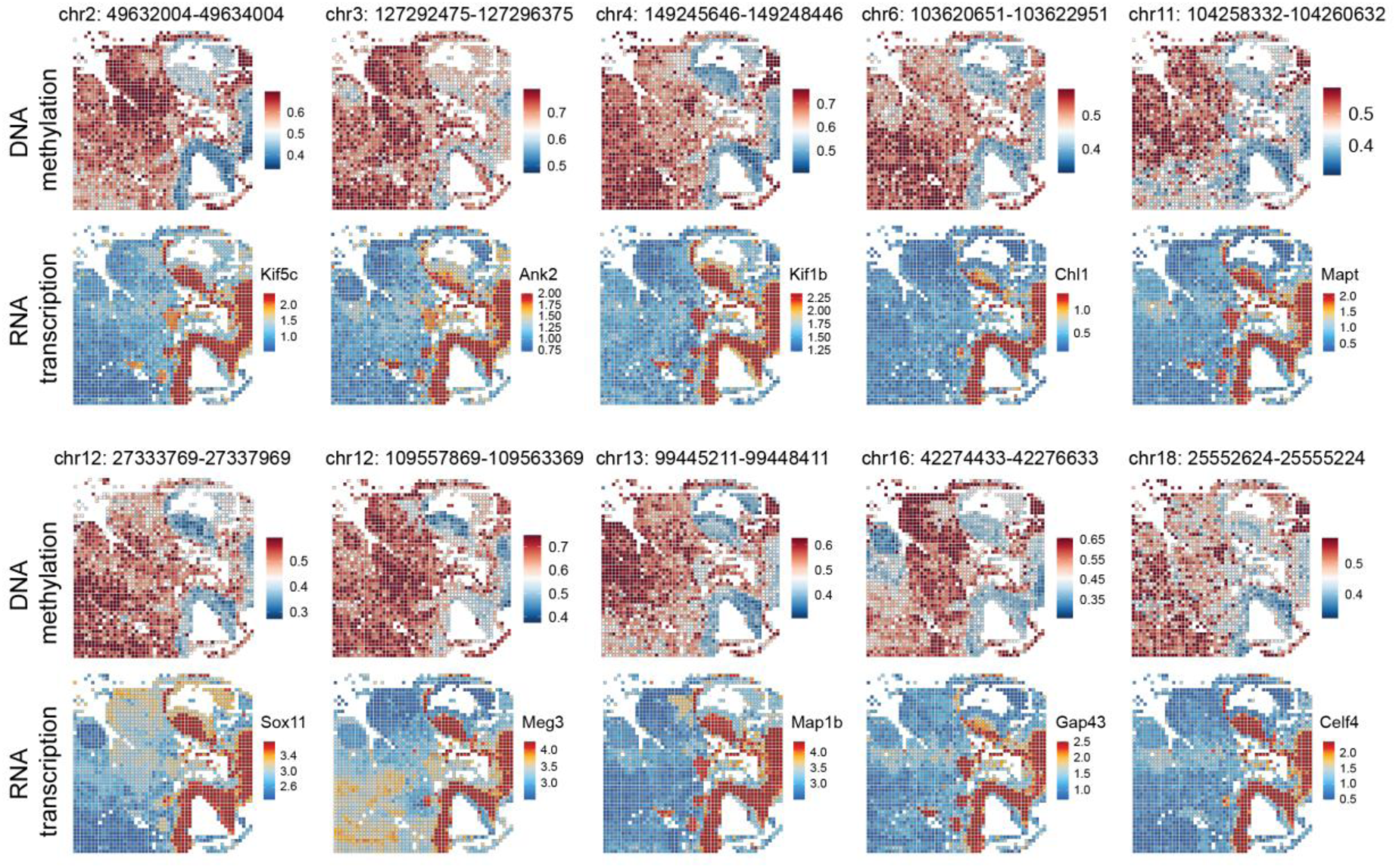
Spatial mapping of DNA methylation and RNA expression in the brain and spinal cord region of E11 mouse embryo. Heatmaps illustrating the spatial distribution of DNA methylation levels (top rows) and RNA expression levels (bottom rows) for marker genomic loci and corresponding genes in the brain and spinal cord regions (W2) of E11 mouse embryo.

**Extended Data Fig. 7:**
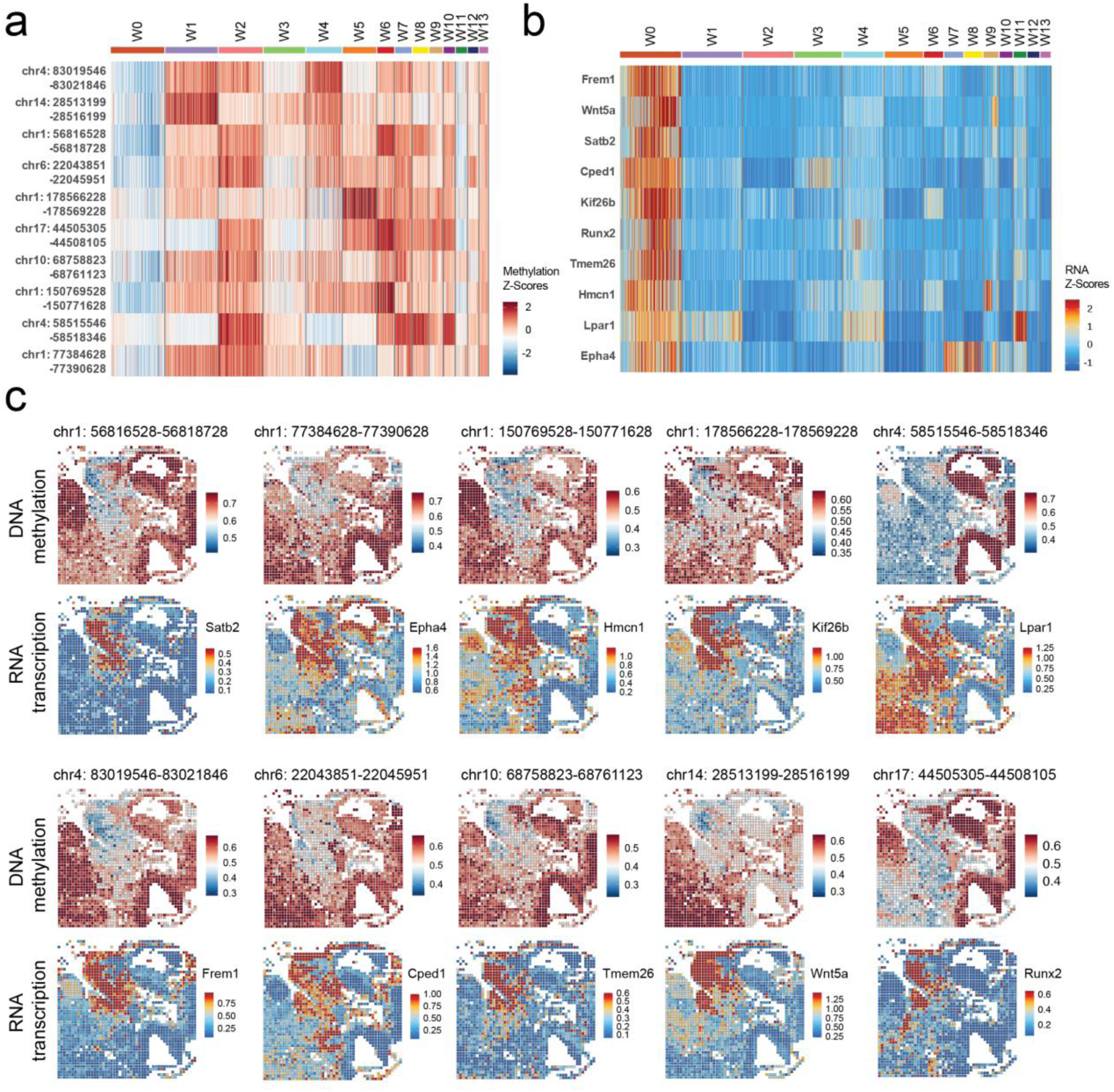
Spatial mapping of DNA methylation and RNA expression in the craniofacial region of E11 mouse embryo. **a,** Heatmap of DNA methylation levels for the top 10 differentially methylated genomic loci in the craniofacial region (cluster W0) of E11 mouse embryo. Each row represents a specific genomic locus, and each column represents a different cluster. The color scale indicates the Z-scores of DNA methylation levels. **b,** Heatmap of expression levels for nearby genes corresponding to the genomic loci in **a**. Each row represents a specific gene, and each column represents a different cluster. The color scale indicates the Z-scores of gene expression levels. **c,** Spatial mapping of DNA methylation and RNA expression levels for selected marker genes in the craniofacial region (W0) of E11 mouse embryo.

**Extended Data Fig. 8:**
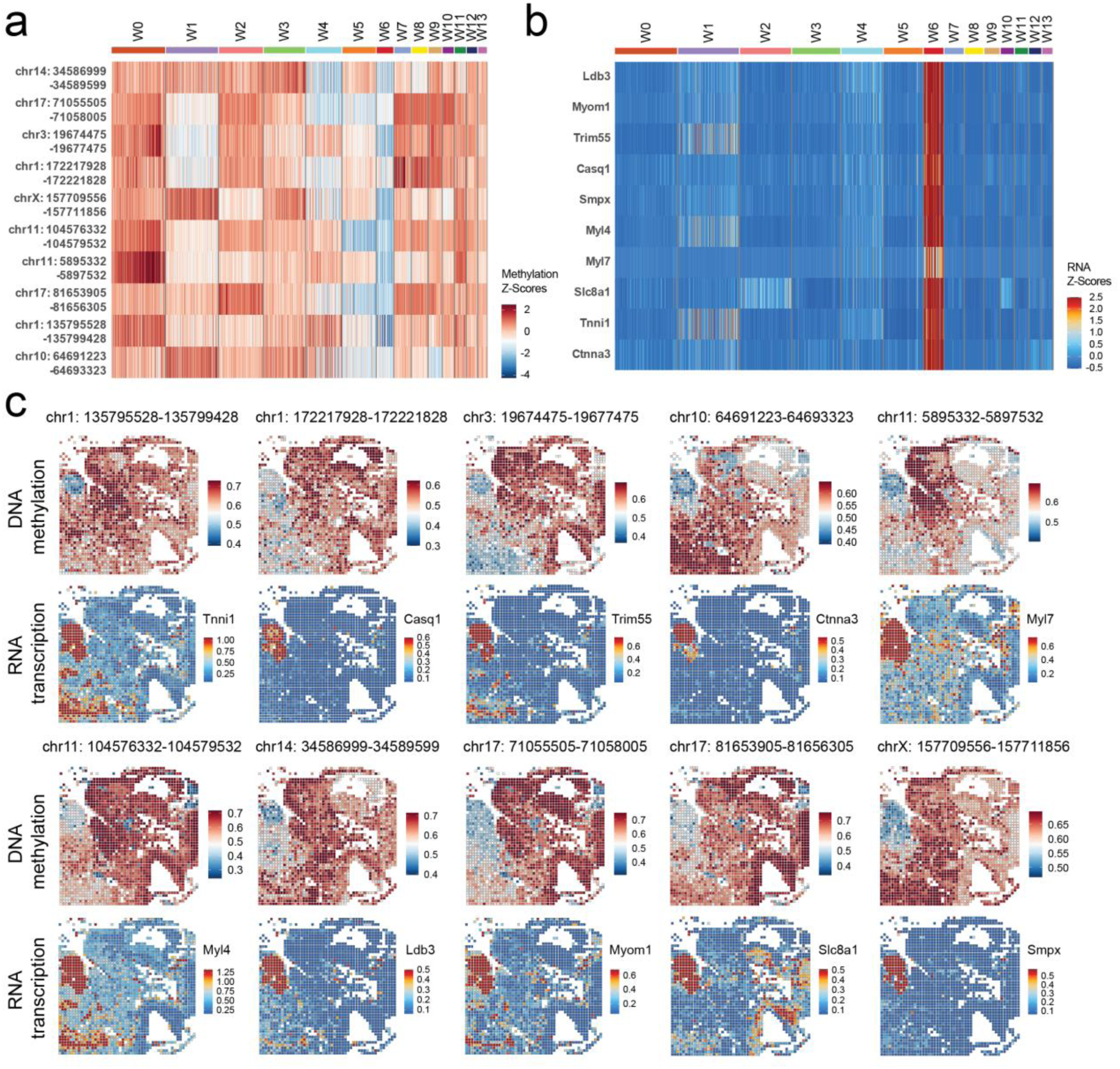
Spatial mapping of DNA methylation and RNA expression in the heart region of E11 mouse embryo. **a,** Heatmap of DNA methylation levels for the top 10 differentially methylated genomic loci in the heart region (cluster W6) of E11 mouse embryo. Each row represents a specific genomic locus, and each column represents a different cluster. The color scale indicates the Z-scores of DNA methylation levels. **b,** Heatmap of RNA expression levels for nearby genes corresponding to the genomic loci in **a**. Each row represents a specific gene, and each column represents a different cluster. The color scale indicates the Z-scores of gene expression levels. **c,** Spatial mapping of DNA methylation and RNA expression levels for selected marker genes in the heart region (W6) of E11 mouse embryo.

**Extended Data Fig. 9:**
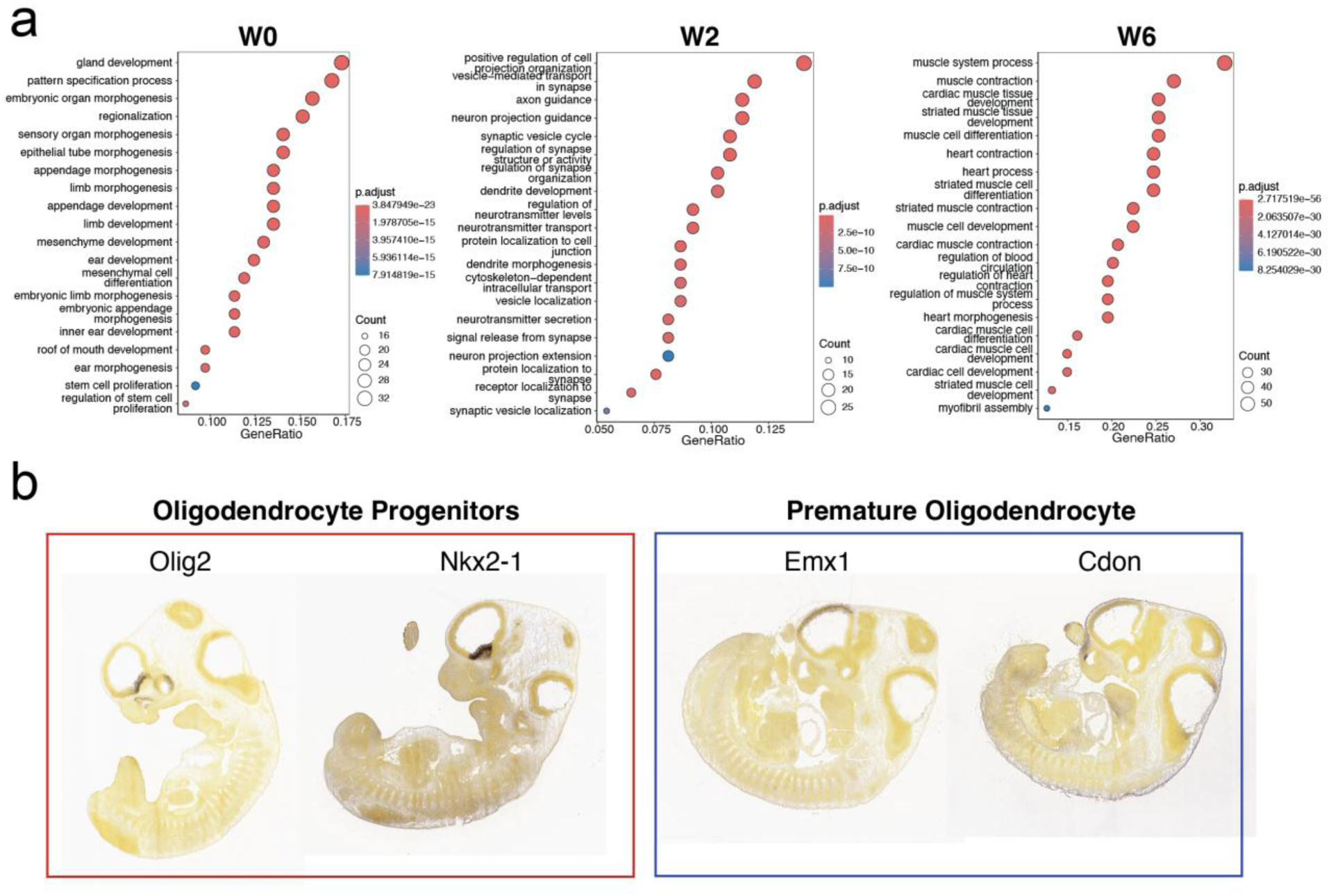
Further analysis of DNA methylation and RNA expression in E11 embryo. **a,** GO enrichment analysis for differentially expressed genes in clusters W0, W2, and W6. The dot plots show the top GO terms for biological processes enriched in each cluster. The x-axis represents the GeneRatio, and the y-axis lists the GO terms. The size of the dots indicates the count of genes associated with each term, and the color represents the adjusted p-value (p.adjust). **b,** In situ hybridization (ISH) images from the Allen Developing Mouse Brain Atlas^84^ show the spatial distribution of markers of oligodendrocyte progenitors and premature oligodendrocytes in the E11 mouse embryo. The images display the expression patterns of markers *Olig2* and *Nkx2-1* for oligodendrocyte progenitors, and *Emx1* and *Cdon* for premature oligodendrocytes. These ISH images highlight the regions where these progenitor and premature cells are localized within the tissue.

**Extended Data Fig. 10:**
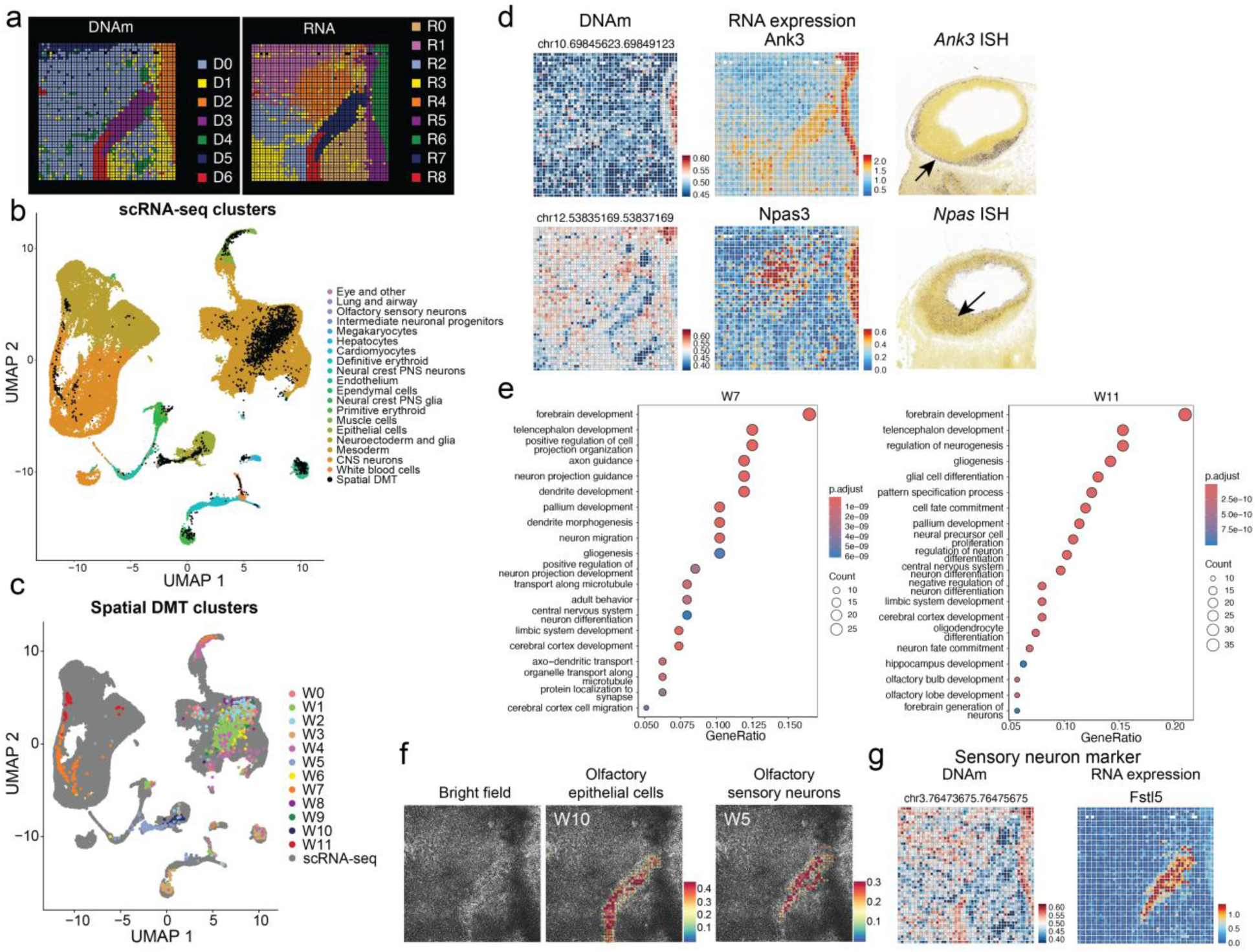
Further analysis of DNA methylation and RNA expression in E11 facial and forebrain regions (pixel size, 10 μm). **a,** Spatial distribution of all clusters for DNA methylation (left) and RNA transcription (right). **b-c,** Integration of Spatial-DMT RNA data with scRNA-seq data from mouse embryo^36^. Cells are colored according to cell annotations from scRNA-seq data^36^ **(b)** and unsupervised clustering results from Spatial-DMT **(c)**. **d,** Spatial mapping of DNA methylation, RNA expression levels, and in situ hybridization (ISH) images from the Allen Developing Mouse Brain Atlas^84^ show the spatial distribution of selected markers in clusters W7 and W11. **e,** GO enrichment analysis for differentially expressed genes in clusters W7 and W11. The dot plots show the top GO terms for biological processes enriched in each cluster. The x-axis represents the GeneRatio, and the y-axis lists the GO terms. The size of the dots indicates the count of genes associated with each term, and the color represents the adjusted p-value (p.adjust). **f,** Spatial distribution of olfactory epithelial cells and olfactory sensory neurons revealed by cell type decomposition of spatial transcriptomic pixels using a scRNA-seq reference^36^. **g,** Spatial mapping of DNA methylation and gene expression level of *Fstl5* (olfactory sensory neurons marker).

**Extended Data Fig. 11:**
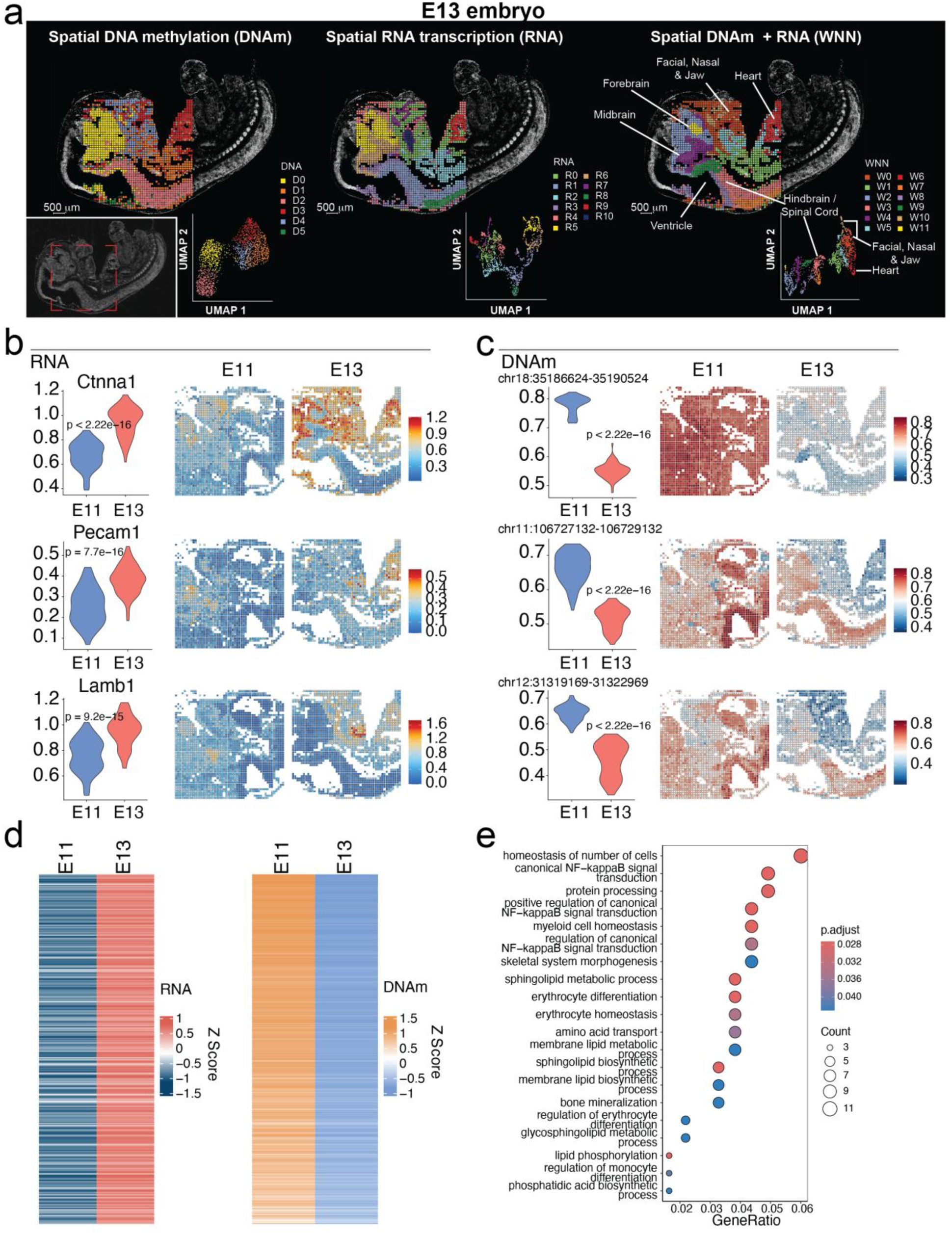
Comparative analysis of DNA methylation and RNA expression between E11 and E13 mouse embryos. **a,** Spatial distribution and UMAP visualization of clusters identified from spatial DNA methylation data (left), spatial RNA transcription data (middle), and integrated DNA and RNA data using WNN analysis (right) for the E13 embryo. Each color represents a different cluster, illustrating the spatial distribution of methylation patterns, gene expression profiles, and the integrated data across different regions such as the forebrain, midbrain, hindbrain/spinal cord, heart, and facial/nasal/jaw region. **b, c,** Comparative analysis of RNA expression **(b)** and DNA methylation levels **(c)** for upregulated genes in E13 heart region, indicating significant changes in methylation and gene expression patterns during embryonic development. **d,** Heatmaps comparing global RNA expression (left) and DNA methylation levels (right) in brain and spinal cord regions between E11 and E13 embryos. **e,** GO enrichment analysis for biological processes related to demethylated and upregulated genes in the heart region of E13 embryo. The size of the dots indicates the count of genes associated with each term, and the color represents the adjusted p-value (p.adjust).

**Extended Data Fig. 12:**
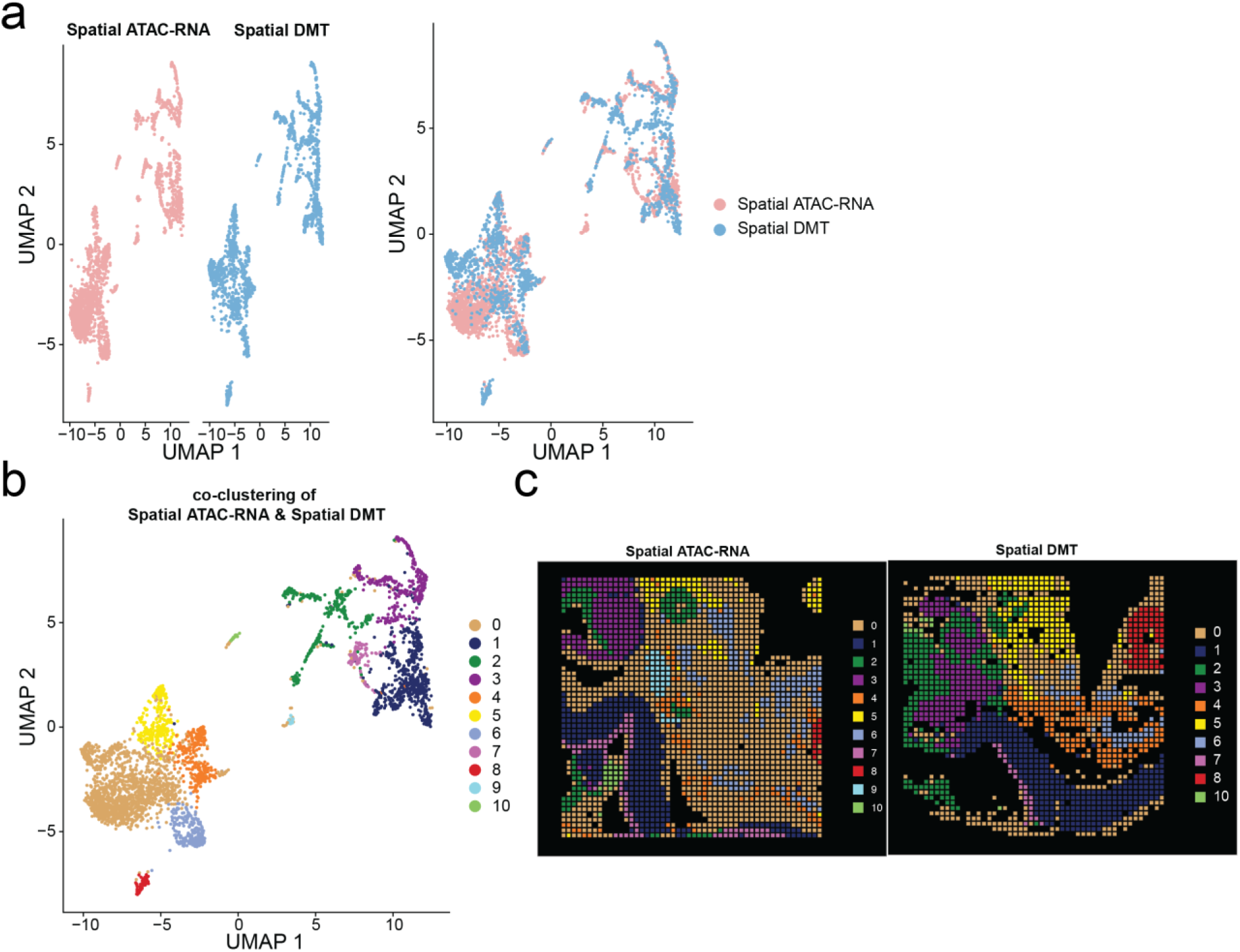
Integrative analysis of E13 embryo data. a-b, Integrated visualization of Spatial-DMT and spatial-ATAC-RNA-seq^18^ datasets from E13 mouse embryos. Cells are colored according to experiments **(a)** and unsupervised co-clustering results **(b)**. **c,** Spatial distribution of co-clustering results, comparing spatial-ATAC-RNA-seq (left) and Spatial-DMT (right). Spatial maps show distinct clustering patterns, with each color representing the shared clusters defined from integrated analysis in **(b)**. Concordant spatial domains were shown across both technologies.

**Extended Data Fig. 13:**
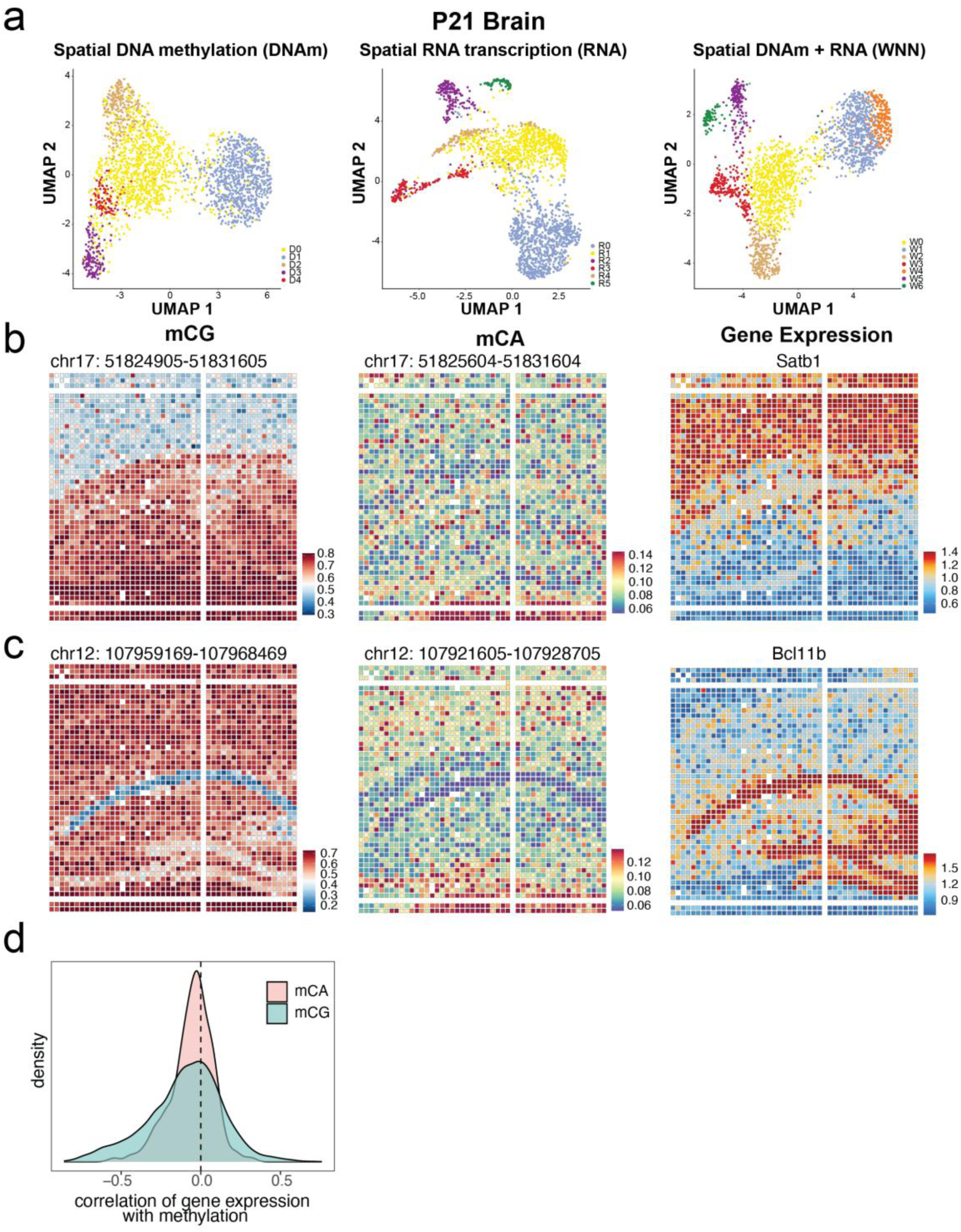
Analysis of DNA methylation and RNA expression in the P21 mouse brain. **a,** UMAP visualization of clusters identified from spatial DNA methylation data (left), spatial RNA transcription data (middle), and integrated DNA and RNA data using WNN analysis (right) for the P21 brain. **b,** Spatial mapping of CpG (left) and CpA (middle) methylation levels, and RNA expression levels (right) for *Satb1* in the P21 brain. **c,** Spatial mapping of CpG (left) and CpA (middle) methylation levels, and RNA expression levels (right) for *Bcl11b* in the P21 brain. **d,** Density plot showing the correlation of gene expression with CpA and CpG methylation in the P21 mouse brain. The plot illustrates the distribution of correlation coefficients, with red indicating CpA methylation and green indicating CpG methylation.

**Extended Data Fig. 14:**
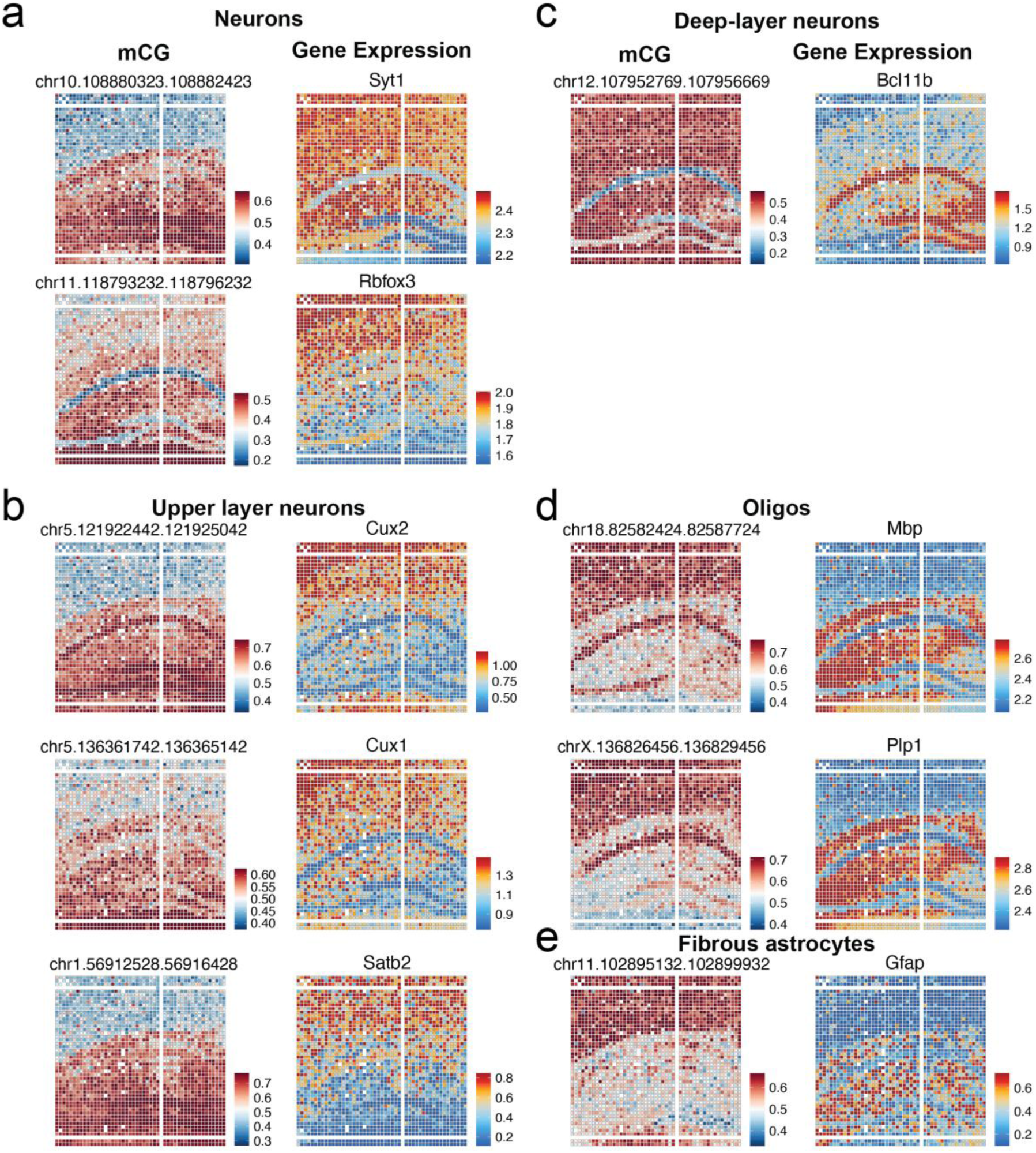
Spatial mapping of DNA methylation and RNA expression levels of selected markers across distinct cell types in P21 mouse brain. Heatmaps illustrating the spatial distribution of DNA methylation levels and RNA expression levels for markers of neurons (*Syt1*, *Rbfox3*) **(a)**, upper layer neurons (*Cux2*, *Cux1*, *Satb2*) **(b)**, deep-layer neurons (*Bcl11b*) **(c)**, oligos (*Mbp*, *Plp1*) **(d)**, and fibrous astrocytes (*Gfap*) **(e)**.

**Extended Data Fig. 15:**
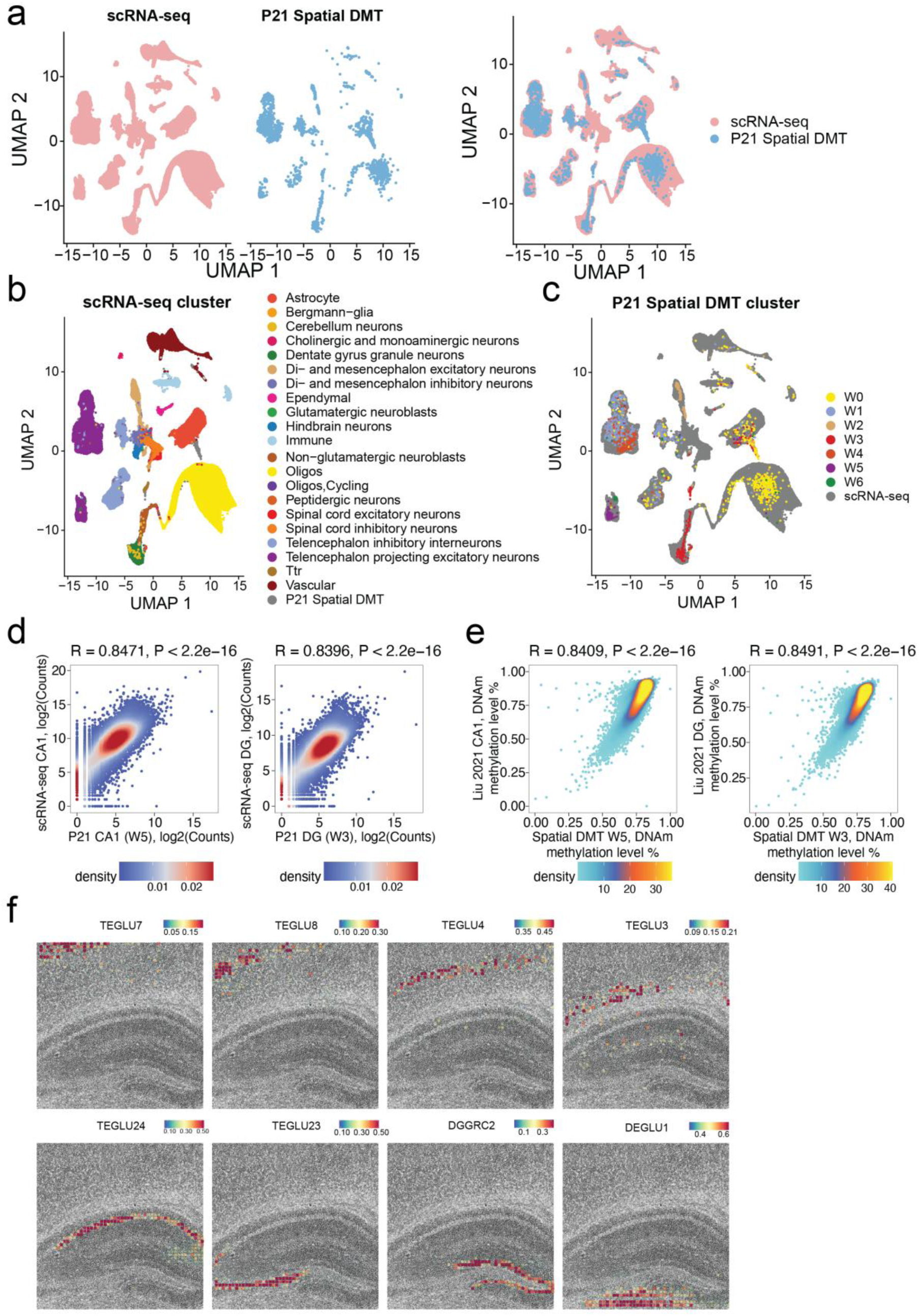
Integrative analysis of P21 mouse brain data. **a-c,** Integrated visualization of Spatial-DMT and scRNA-seq datasets from adolescent mouse brain^61^. Cells are colored according to experiments **(a)**, cell annotations from scRNA-seq data^61^ **(b)**, and unsupervised clustering results from Spatial-DMT **(c)**. **d,** Correlation of gene expression profiles from cluster W5 (CA1) and W3 (DG) with a reference scRNA-seq dataset^61^. **e,** Correlation of DNA methylation profiles (100kb bin) from cluster W5 (CA1) and W3 (DG) with a single-cell methylation dataset^22^. **f,** Spatial distribution of selected cell types resolved by cell-type decomposition of spatial transcriptomic pixels using a reference scRNA-seq dataset^61^, including cerebral cortex excitatory neurons (TEGLU7, TEGLU8, TEGLU4, TEGLU3), cornu ammonis excitatory neurons (TEGLU24 and TEGLU23), dentate gyrus granule neurons (DGGRC2), and thalamus excitatory neurons (DEGLU1).

**Extended Data Fig. 16:**
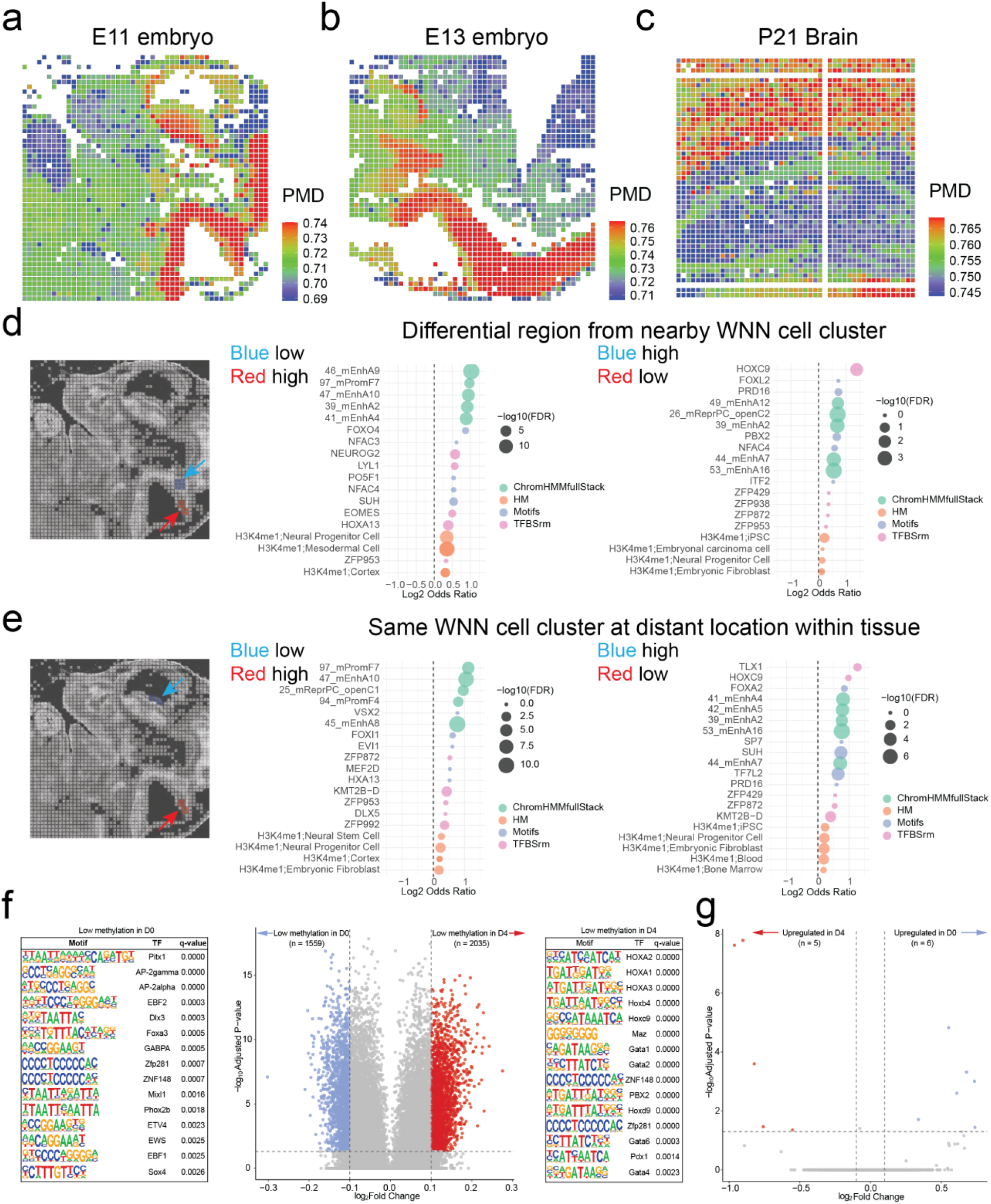
Spatial-DMT resolves mitotic history and subtle region-specific epigenetic variations. **a–c**, Methylation levels at Partially Methylated Domains (PMDs) in the E11 mouse embryo (**a**), E13 mouse embryo (**b**), and P21 mouse brain (**c**). **d**, Differential region analysis between adjacent WNN cell clusters. The middle panel shows chromatin features enriched at hypomethylated genome loci in the blue region, while the right panel corresponds to those enriched in the red region. A summary of ChromHMMfullStack annotation is provided in Supplementary Table 6. **e,** Differential region analysis within the same WNN cell cluster but across spatially distant locations. Chromatin features enriched at hypomethylated loci in the blue and red regions are shown in the middle and right panels, respectively. FDR, False Discovery Rate. **f,** Enriched TF motifs in differentially hypomethylated VMRs between D0 and D4 cluster pixels derived from R3 cluster in the E11 embryo. **g,** Differentially expressed genes corresponding to the D0 and D4 pixels originated from the R3 cluster in the E11 embryo.

